# Thermodynamics shape the *in vivo* enzyme burden of glycolytic pathways

**DOI:** 10.1101/2025.01.31.635972

**Authors:** Daven B. Khana, Annie Jen, Evgenia Shishkova, Eashant Thusoo, Jonathan Williams, Alex Henkel, David M. Stevenson, Joshua J. Coon, Daniel Amador-Noguez

## Abstract

Thermodynamically constrained reactions and pathways are hypothesized to impose greater protein demands on cells, requiring higher enzyme amounts to sustain a given flux compared to those with stronger thermodynamics. To test this, we quantified the absolute concentrations of glycolytic enzymes in three bacterial species —*Zymomonas mobilis*, *Escherichia coli*, and *Clostridium thermocellum*— which employ distinct glycolytic pathways with varying thermodynamic driving forces.

By integrating enzyme concentration data with corresponding *in vivo* metabolic fluxes and *ΔG* measurements, we found that the highly favorable Entner-Doudoroff (ED) pathway in *Z. mobilis* requires only one-fourth the amount of enzymatic protein to sustain the same flux as the thermodynamically constrained pyrophosphate-dependent glycolytic pathway in *C. thermocellum*, with the Embden-Meyerhof-Parnas (EMP) pathway in *E. coli* exhibiting intermediate thermodynamic favorability and enzyme demand. Across all three pathways, early reactions with stronger thermodynamic driving forces generally required lower enzyme investment than later, less favorable steps. Additionally, reflecting differences in glycolytic strategies, the highly reversible ethanol fermentation pathway in *C. thermocellum* requires 10-fold more protein to maintain the same flux as the irreversible, forward-driven ethanol fermentation pathway in *Z. mobilis*.

Thus, thermodynamic driving forces constitute a major *in vivo* determinant of the enzyme burden in metabolic pathways.

## 1. Introduction

Metabolic flux is a primary driver of cellular physiology. Cells regulate fluxes to meet energy and biosynthetic demands while efficiently managing limited resources, including the finite capacity to synthesize and maintain metabolic enzymes^1–4^. Multiple factors influence metabolic flux within cells, including enzyme abundance, catalytic efficiency (k_cat_), active site saturation (governed by K_m_ values and substrate concentrations), and regulatory mechanisms such as allosteric inhibition and post-translational modifications^5–10^. A less commonly appreciated but critical factor is the energetics of biochemical reactions, typically quantified as the change in Gibbs free energy (*ΔG*). This thermodynamic parameter not only determines reaction directionality but also imposes intrinsic constraints on flux^11,12^. Specifically, the ratio of forward (*J^+^*) to reverse (*J^−^*) fluxes of a reaction relates to its *ΔG* via the equation:

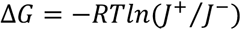

where *R* is the gas constant, and *T* is the absolute temperature in kelvin. This equation, known as the flux-force relationship, reveals the interdependence between a reaction’s thermodynamic driving force, net flux, and enzyme cost^5,13–15^. Reactions far from thermodynamic equilibrium (i.e., with a large negative *ΔG*) have forward fluxes that greatly exceed reverse fluxes, resulting in a high net flux (*J^net^*= J^+^ − J^−^) and efficient enzyme utilization, as most enzyme activity is directed toward the forward reaction. In contrast, reactions operating near equilibrium have nearly equal forward and reverse fluxes (*J⁺ ≈ J⁻*), which leads to inefficient enzyme utilization and a reduced net flux. Consequently, thermodynamically constrained reactions incur higher enzyme costs —defined as the amount of enzyme required per unit flux— to sustain the same net flux compared to reactions with stronger thermodynamic driving forces^5,16^.

Building on these principles, a previous computational study investigated the interdependence between pathway thermodynamics, enzyme cost, and energy output (i.e., ATP production) in the two most prevalent glycolytic pathways used by bacteria: the Embden-Meyerhof-Parnas (EMP) and the Entner-Doudoroff (ED) pathways^17^. By combining computationally estimated free energies with model-derived protein cost estimates, this study showed that the ED pathway is significantly less thermodynamically constrained than the EMP pathway and predicted that the ED pathway requires three to five times less enzymatic protein to sustain the same glycolytic flux as the EMP pathway. However, this reduction in enzyme cost, driven by greater thermodynamic favorability, comes at the expense of a lower ATP yield per glucose^17^. Subsequent computational studies have further supported the hypothesis that thermodynamically constrained reactions and pathways impose greater protein demands on the cell as a consequence of large reverse fluxes and inefficient enzyme utilization^10,16,18^.

While these computational predictions are compelling, they remain to be experimentally validated. Testing these hypotheses *in vivo* requires simultaneous measurements of metabolic fluxes and protein levels in organisms that utilize pathways with distinct thermodynamic profiles^17^. In this study, we address this gap by quantifying the absolute concentrations of glycolytic enzymes in three bacterial species −*Zymomonas mobilis*, *Escherichia coli*, and *Clostridium thermocellum*− which employ distinct glycolytic pathways with varying thermodynamic driving forces. By integrating enzyme concentration data with corresponding *in vivo* metabolic fluxes and intracellular *ΔG* measurements, we provide strong experimental evidence that thermodynamic driving forces play a crucial role in determining the *in vivo* enzyme burden of metabolic reactions and pathways.

## 2. Results

### 2.1 Experimental system: energetics and flux of three distinct glycolytic pathways

We investigated the *in vivo* relationship between pathway thermodynamics, metabolic fluxes, and enzyme concentrations across the glycolytic pathways of three different bacteria: the ethanologenic *Z. mobilis*, the cellulolytic and ethanologenic *C. thermocellum*, and the model organism *E. coli*. These bacteria metabolize glucose to pyruvate via distinct glycolytic routes, which vary in key enzymatic steps, energy yield (i.e., ATP/ GTP output), thermodynamics, and flux (Figure 1). *Z. mobilis* exclusively relies on the ED pathway for glucose catabolism (Figure 1A)^19–21^. In contrast, *C. thermocellum* employs a pyrophosphate (PPi)-dependent EMP pathway (PPi-EMP), which features a PPi-phosphofructokinase (PPi-Pfk) that utilizes PPi, rather than ATP, as a phosphate donor to convert fructose 6-phosphate (F6P) to fructose 1,6-bisphosphate (FBP)^22–25^. Additionally, *C. thermocellum* lacks a pyruvate kinase (Pyk) to convert phosphoenolpyruvate (PEP) to pyruvate. Instead, it produces pyruvate via a PPi-dependent dikinase (Ppdk), and can also generate pyruvate via the ‘malate shunt’, which involves phosphoenolpyruvate carboxykinase (Pepck), malate dehydrogenase (Mdh), and malic enzyme (Me) (Figure 1A)^22,26,27^. Finally, *E. coli* primarily uses the EMP pathway to convert glucose into pyruvate, utilizing the ED pathway only under specific conditions, such as growth on gluconate or during gut colonization^28,29^. A notable difference between *E. coli*, *Z. mobilis*, and *C. thermocellum* lies in glucose uptake and its conversion to glucose 6-phosphate (G6P). In *E. coli*, glucose import into the cytoplasm is coupled with its phosphorylation to G6P via the phosphotransferase system (PTS), which uses PEP as the phosphate donor and produces pyruvate as a byproduct. In contrast, *Z. mobilis* and *C. thermocellum* phosphorylate glucose or cellobiose, respectively, only after these sugars enter the cytoplasm.

**Figure 1.**
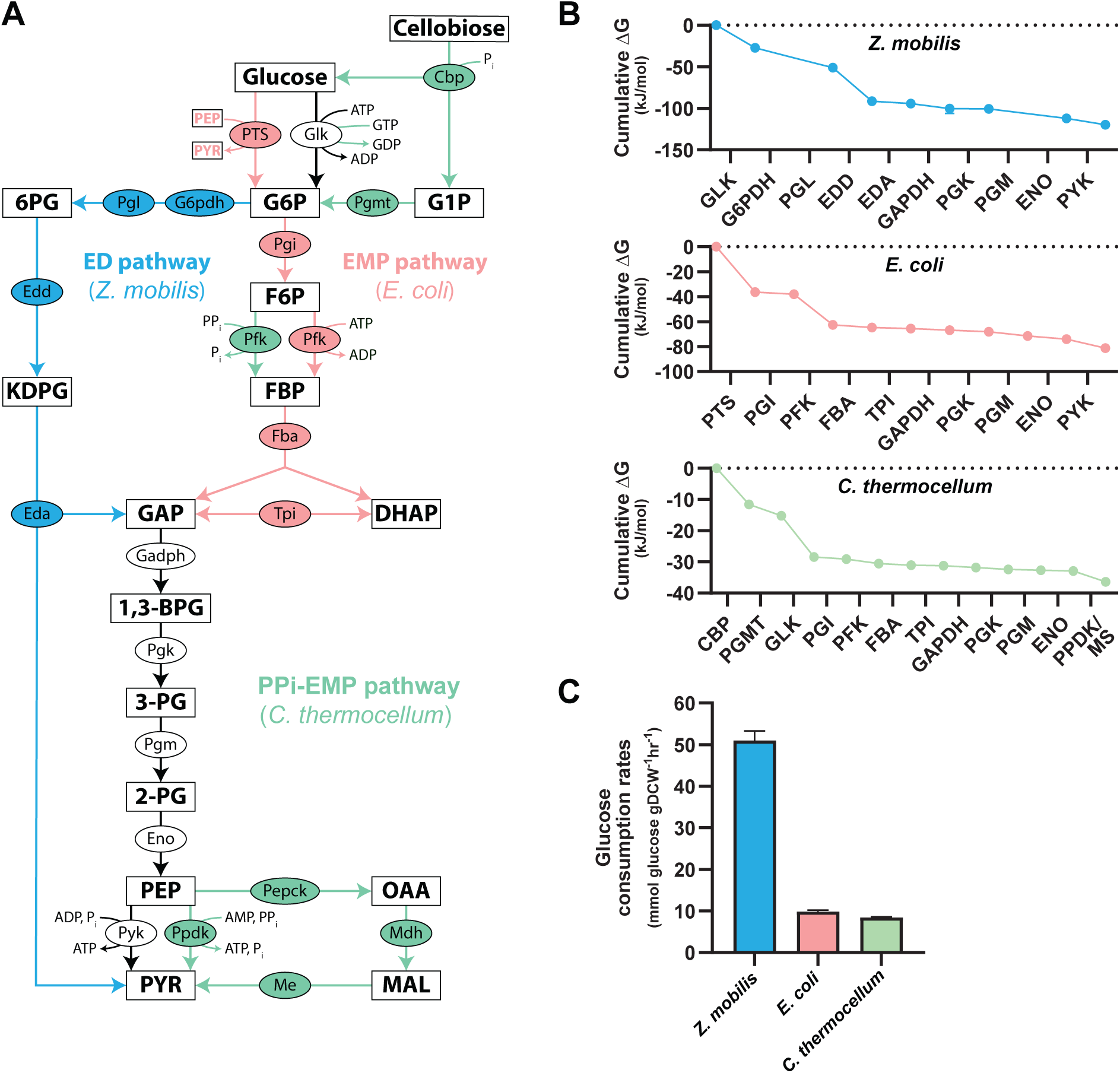
Glycolytic pathways and their energetics. **A.** The Entner-Doudoroff (ED) pathway in *Z. mobilis* (blue arrows), the Embden–Meyerhof–Parnas (EMP) pathway in *E. coli* (pink arrows), and the PPi-EMP pathway in *C. thermocellum* (green arrows) utilize distinct enzymes at various steps to convert glucose into pyruvate (PYR). Reactions depicted with black arrows are common to all three pathways. In *E. coli*, glucose is simultaneously imported and converted to glucose 6-phosphate (G6P) using phosphoenolpyruvate (PEP) as the phosphate donor via the phosphotransferase system (PTS). Enzymes are depicted as ovals, and metabolites are shown as rectangles. **B.** The cumulative drop in *ΔG* for the glycolytic pathways in *Z. mobilis* (blue), *E. coli* (pink), and *C. thermocellum* (green). *ΔG* data are a combination of previous experimental measurements^30–32^ and computationally estimated values, constrained by *in vivo* metabolite concentrations, obtained in this work (Materials and Methods). The *ΔG* for the pyrophosphate dependent pyruvate dikinase (PPDK) and malate shunt (MS) (i.e., PEP carboxykinase (PEPCK), malate (MAL) dehydrogenase (MDH), malic enzyme (ME)) in *C. thermocellum* represents the combined reaction (Table S1, Materials and Methods). **C.** Glucose consumption rates for each bacterium. The glucose consumption rate for *C. thermocellum* is presented as twice the calculated cellobiose uptake rate, since each molecule of cellobiose consists of two glucose moieties. Glucose uptake rates were calculated in cells grown aerobically (*E. coli*) or anaerobically (*Z. mobilis* and *C. thermocellum*) in minimal media (Materials and Methods). Data represent the averages of 3-4 biological replicates. Error bars show 95% confidence intervals (*ΔG* values) or ± standard deviation (sugar consumption rates). Some error bars are too small to be visible in this representation. See Table S1 and Table S2 for *ΔG* and glucose consumption rate data, respectively. Metabolite abbreviations: 6-phosphogluconate (6PG), glucose 6-phosphate (G6P), glucose 1-phosphate (G1P), 2-keto-3-deoxy-6-phosphogluconate (KDPG), fructose 6-phosphate (F6P), fructose 1,6-bisphosphate (FBP), glyceraldehyde 3-phosphate (GAP), dihydroxyacetone phosphate (DHAP), 1,3-bisphosphoglycerate (1,3-BPG), 3-phosphoglycerate (3-PG), 2-phosphoglycerate (2-PG), phosphoenolpyruvate (PEP), pyruvate (PYR), oxaloacetate (OAA), malate (MAL). Enzyme abbreviations: cellobiose phosphorylase (Cbp), glucokinase (Glk), G6P dehydrogenase (G6pdh), phosphogluconolactonase (Pgl), phosphoglucomutase (Pgmt), 6PG dehydratase (Edd), phosphoglucose isomerase (Pgi), phosphofructokinase (Pfk), KDPG aldolase (Eda), FBP aldolase (Fba), triose phosphate isomerase (Tpi), GAP dehydrogenase (Gapdh), phosphoglycerate kinase (Pgk), phosphoglycerate mutase (Pgm), enolase (Eno), PYR kinase (Pyk), PYR phosphate dikinase (Ppdk), PEP carboxykinase (Pepck), MAL dehydrogenase (Mdh), malic enzyme (Me).

The overall thermodynamic favorability and energy output of these glycolytic pathways differ greatly. *In vivo ΔG* measurements obtained from ^13^C and ^2^H metabolic flux analyses (MFA)^30–32^ coupled with *ΔG* computational estimates (Materials and Methods) show that the ED pathway in *Z. mobilis* is approximately three times more thermodynamically favorable than the PPi-EMP pathway in *C. thermocellum* and nearly twice as favorable as the EMP pathway in *E. coli* (Figure 1B, Table S1). Notably, the high thermodynamic favorability of the ED pathway in *Z. mobilis* correlates with an *in vivo* glycolytic rate that is approximately 6-fold higher than that of *C. thermocellum* and 5-fold higher than that of *E. coli* (Figure 1C, Table S2).

### 2.2 Protein resources are unevenly allocated across glycolysis

In *Z. mobilis*, each reaction of the Entner-Doudoroff (ED) glycolytic pathway is catalyzed by a single enzyme^33^. In contrast, *E. coli* has multiple isoenzymes for several glycolytic reactions, including phosphofructokinase (PFK), fructose 1,6-bisphosphate aldolase (FBA), glyceraldehyde 3-phosphate dehydrogenase (GAPDH), phosphoglycerate mutase (PGM), and pyruvate kinase (PYK) ^34–42^. Similarly, *C. thermocellum* possesses multiple isoenzymes for FBA and PGM (Table S3)^43^.

We used shotgun proteomics to identify the predominant glycolytic enzymes in each bacterium (Table S3). *Z. mobilis* and *C. thermocellum* were grown anaerobically, while *E. coli* was cultured under aerobic conditions. *Z. mobilis* and *E. coli* were grown using glucose as the sole carbon source, whereas *C. thermocellum* was grown on cellobiose (Materials and Methods). All isoenzymes with comparable expression levels, as determined by intensity-based absolute quantification (iBAQ) values^44,45^ from shotgun proteomics, were selected for direct quantitation using the absolute quantification (AQUA) method (Table S3). For each protein, 2 to 8 isotopically labeled reference peptides were chosen based on shotgun proteomics data (Table S5)^49,50^. Isoenzymes with markedly lower expression (e.g., >15-fold difference) compared to the predominant isoenzyme were excluded from AQUA quantification (Table S3).

ED pathway enzymes in *E. coli* were also excluded from direct absolute quantitation using AQUA as previous MFA studies have shown negligible carbon flux (0.2-1%) through 6-phosphogluconate dehydratase (EDD) and 2-dehydro-3-deoxyphosphogluconate aldolase (EDA) when *E. coli* is grown aerobically on glucose^46,47^. Similarly, although *C. thermocellum* possesses multiple ATP/GTP dependent PFKs in addition to PPi-Pfk^43^, enzyme assays in cell extracts revealed no ATP/GTP-PFK activity^22,48^. Consistent with these findings and other previous studies^22–24^, PPi-PFK was the most highly expressed PFK isozyme in our *C. thermocellum* cells, leading us to exclude ATP/GTP-dependent PFKs from direct AQUA quantitation (Table S3).

Using AQUA, we determined the absolute intracellular concentrations of 13, 16, and 15 glycolytic enzymes in *Z. mobilis*, *C. thermocellum*, and *E. coli*, respectively (Table 1, Table S4). For *Z. mobilis* and *C. thermocellum*, both of which produce ethanol as their primary fermentation product, we also used AQUA to quantify the absolute concentrations of their ethanol pathway enzymes (Table 1, Table S4).

**Table 1.** Absolute intracellular concentrations of glycolytic and fermentation enzymes in *Z. mobilis*, *C. thermocellum*, and *E. coli* quantified via AQUA.

| Organism | Locus Tag | Protein | Abbr. | Avg.<br>(fg/cell) | SD |
| --- | --- | --- | --- | --- | --- |
| *Z. mobilis* | ZMO0366 | Glucose facilitated diffusion protein | GlF | 0.403 | 0.016 |
| *Z. mobilis* | ZMO0369 | Glucokinase | GlK | 0.498 | 0.068 |
| *Z. mobilis* | ZMO0367 | Glucose 6-phosphahte dehydrogenase | G6pdh | 1.417 | 0.366 |
| *Z. mobilis* | ZMO1478 | 6-phosphogluconolactonase | Pgl | 0.111 | 0.037 |
| *Z. mobilis* | ZMO0368 | 6-phosphogluconate dehydratase | Edd | 1.625 | 0.144 |
| *Z. mobilis* | ZMO0997 | 2-dehydro-3-deoxyphosphogluconate aldolase | Eda | 0.611 | 0.300 |
| *Z. mobilis* | ZMO0177 | Glyceraldehyde 3-phosphate dehydrogenase | Gapdh | 4.498 | 0.840 |
| *Z. mobilis* | ZMO0178 | Phosphoglycerate kinase | Pgk | 1.465 | 0.200 |
| *Z. mobilis* | ZMO1240 | Phosphoglycerate mutase | Pgm | 1.003 | 0.059 |
| *Z. mobilis* | ZMO1608 | Enolase | Eno | 2.899 | 0.110 |
| *Z. mobilis* | ZMO0152 | Pyruvate kinase | Pyk | 4.338 | 0.273 |
| *Z. mobilis* | ZMO1212 | Phosphoglucose isomerase | Pgi | 0.449 | 0.175 |
| *Z. mobilis* | ZMO0179 | Fructose biphosphate aldolase | Fba | 0.048 | 0.018 |
| *Z. mobilis* | ZMO0465 | Triose phosphate isomerase | Tpi | 0.033 | 0.014 |
| *Z. mobilis* | ZMO1360 | Pyruvate decarboxylase | Pdc | 6.662 | 0.477 |
| *Z. mobilis* | ZMO1236 | Alcohol dehydrogenase I | AdhA | 0.067 | 0.013 |
| *Z. mobilis* | ZMO1596 | Alcohol dehydrogenase II | AdhB | 1.643 | 0.642 |
| *E. coli* | b1101 | Phosphotransferase enzyme IIBC component | PtsG | 0.264 | 0.057 |
| *E. coli* | b2388 | Glucokinase | GlK | 0.043 | 0.008 |
| *E. coli* | b4025 | Phosphoglucose isomerase | Pgi | 0.216 | 0.032 |
| *E. coli* | b3916 | 6-phosphofructokinase I | PfkA | 0.112 | 0.009 |
| *E. coli* | b1723 | 6-phosphofructokinase II | PfkB | 0.026 | 0.009 |
| *E. coli* | b2097 | Fructose biphosphate aldolase class I | FbaB | 0.042 | 0.008 |
| *E. coli* | b2925 | Fructose biphosphate aldolase class II | FbaA | 0.227 | 0.093 |
| *E. coli* | b3919 | Triose phosphate isomerase | Tpi | 0.207 | 0.018 |
| *E. coli* | b1779 | Glyceraldehyde 3-phosphate dehydrogenase | Gapdh | 1.946 | 0.616 |
| *E. coli* | b2926 | Phosphoglycerate kinase | Pgk | 0.831 | 0.054 |
| *E. coli* | b3612 | Phosphoglycerate mutase, 2,3-bisphosphoglycerate independent | GpmM | 0.248 | 0.030 |
| *E. coli* | b0755 | Phosphoglycerate mutase, 2,3-bisphosphoglycerate dependent | GpmA | 0.219 | 0.084 |
| *E. coli* | b2779 | Enolase | Eno | 1.072 | 0.088 |
| *E. coli* | b1676 | Pyruvate kinase I | PykF | 0.354 | 0.033 |
| *E. coli* | b1854 | Pyruvate kinase II | PykA | 0.069 | 0.009 |
| *C. thermocellum* | Clo1313_1954 | Cellobiose phosphorylase | Cbp | 0.254 | 0.052 |
| *C. thermocellum* | Clo1313_0993 | Phosphoglucomutase | Pgmt | 0.088 | 0.016 |
| *C. thermocellum* | Clo1313_0489 | Glucokinase | GlK | 0.057 | 0.010 |
| *C. thermocellum* | Clo1313_2015 | Phosphoglucose isomerase | Pgi | 0.226 | 0.056 |
| *C. thermocellum* | Clo1313_1876 | 6-phosphofructokinase, PPI dependent | Pfk1 | 1.333 | 0.431 |
| *C. thermocellum* | Clo1313_1875 | Fructose bisphosphate aldolase class II | Fba1 | 0.381 | 0.055 |
| *C. thermocellum* | Clo1313_2093 | Triose phosphate isomerase | Tpi | 0.505 | 0.047 |
| *C. thermocellum* | Clo1313_2095 | Glyceraldehyde 3-phosphate dehydrogenase | Gapdh | 2.133 | 0.445 |
| *C. thermocellum* | Clo1313_2094 | Phosphoglycerate kinase | Pgk | 1.285 | 0.207 |
| *C. thermocellum* | Clo1313_2092 | Phosphoglycerate mutase, 2,3-bisphosphoglycerate independent | Pgm1 | 0.217 | 0.046 |
| *C. thermocellum* | Clo1313_0966 | Phosphoglycerate mutase, 2,3-bisphosphoglycerate independent | Pgm2 | 0.018 | 0.003 |
| *C. thermocellum* | Clo1313_2090 | Enolase | Eno | 0.917 | 0.476 |
| *C. thermocellum* | Clo1313_0949 | Pyruvate phosphate dikinase | Ppdk | 0.308 | 0.027 |
| *C. thermocellum* | Clo1313_0415 | Phosphoenolpyruvate carboxykinase | Pepck | 1.206 | 0.477 |
| *C. thermocellum* | Clo1313_1878 | Malate dehydrogenase | Mdh | 0.491 | 0.100 |
| *C. thermocellum* | Clo1313_1879 | Malic Enzyme | Me | 0.542 | 0.182 |
| *C. thermocellum* | Clo1313_0022 | Pyruvate ferredoxin oxidoreductase I, alpha domain | Pfor1-α | 0.676 | 0.044 |
| *C. thermocellum* | Clo1313_1798 | Bifunctional acetaldehyde and alcohol dehydrogenase | Aldh/<br>Adh | 2.244 | 0.299 |

To estimate the absolute concentrations of proteins not quantified via AQUA, we developed a quantitative model based on AQUA-derived absolute protein measurements and their corresponding summed precursor MS intensities (iBAQ values)^44,45^. This approach yielded strong correlations (R² ≈ 0.87–0.94) between AQUA absolute protein measurements and their respective iBAQ values across all three bacteria (Figure S1), with low normalized root mean square error (NRMSE) (Table S4), as determined via leave-one-out-cross-validation (Materials and Methods)^51–53^. Using this method, we quantified the absolute concentrations of 1634, 2428, and 1972 proteins, representing 85%, 56%, and 66% of the proteomes of *Z. mobilis*, *E. coli*, and *C. thermocellum*, respectively (Tables S6-S8).

Figure 2 presents the absolute concentrations of glycolytic enzymes for each bacterium. *Z. mobilis* has approximately three times more total glycolytic enzyme per cell than *E. coli* and twice as much as *C. thermocellum*. Across all three bacteria, protein resources were unevenly distributed within glycolysis, with substantial variation in enzyme abundance at different pathway steps. In *Z. mobilis*, enzymes catalyzing the upper ED pathway (i.e. reactions from glucose to GAP: GLK, G6PDH, PGL, EDD, and EDA) account for only 23% of the total glycolytic enzyme pool on a mass basis (fg cell⁻¹), while enzymes in the lower part of the pathway (i.e. reactions from GAP to pyruvate: GAPDH, PGK, PGM, ENO, and PYK) make up the remaining 77%. A similar trend is observed in *E. coli*, where upper glycolytic enzymes (i.e., GLK, PGI, PFK [PFKA, PFKB], FBA [FBAA, FBAB], TPI) constitute just 16% of the glycolytic enzyme pool, excluding the phosphotransferase system (PTS). Notably, the PTS enzymes themselves represent a major protein investment, comprising 16% of *E. coli*’s glycolytic enzyme pool. In *C. thermocellum*, the enzymes performing the lower glycolytic reactions (i.e., GAPDH, PGK, PGM (PGM1, PGM2), ENO, PPDK, PEPCK, MDH, ME) make up a disproportionate 71% of the total glycolytic enzyme pool. Remarkably, Pfk in upper glycolysis accounts for a much larger fraction of the glycolytic enzyme pool in *C. thermocellum* (13%) compared to *E. coli* (2%).

**Figure 2.**
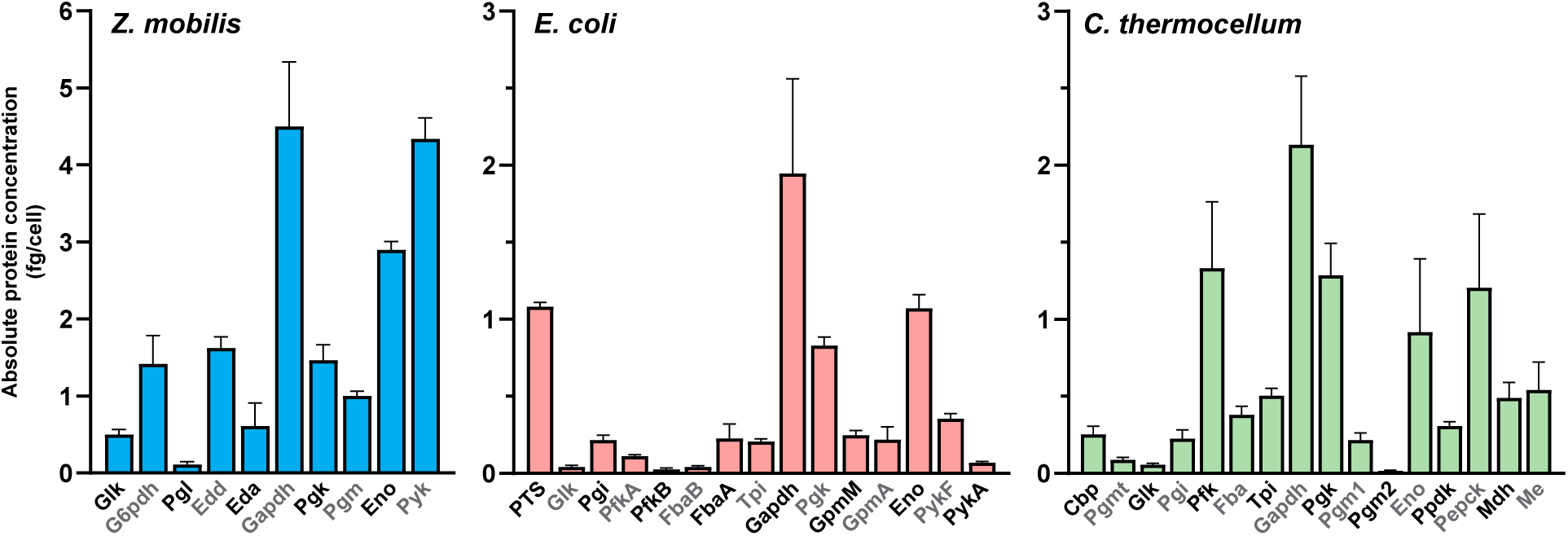
**Intracellular concentrations of glycolytic enzymes.** Absolute concentrations of glycolytic enzymes, expressed in fg per cell, were quantified in *Z. mobilis* (blue), *E. coli* (pink), and *C. thermocellum* (green). The concentration of the phosphotransferase system (PTS) in *E. coli* represents the sum of the four enzymes that comprise the PTS: PtsG, PtsH, PtsI, and Crr (Table S10). Data represent the average of four biological replicates. Error bars show ± standard deviation. Some error bars are too small to be visible in this representation. See Table S4 for absolute enzyme concentration data. Abbreviations: glucokinase (Glk), glucose 6-phosphate dehydrogenase (G6pdh), phosphogluconolactonase (Pgl), 6-phosphogluconate dehydratase (Edd), 2-keto-3-deoxy-6-phosphogluconate aldolase (Eda), glyceraldehyde 3-phosphate dehydrogenase (Gapdh), phosphoglycerate kinase (Pgk), phosphoglycerate mutase (Pgm, GpmM, GpmA, Pgm1, Pgm2), enolase (Eno), pyruvate kinase (Pyk, PykF, PykA), phosphoglucose isomerase (Pgi), fructose 1,6-bisphosphate aldolase (Fba, FbaB, FbaA), triose phosphate isomerase (Tpi), phosphofructokinase (PfkA, PfkB, Pfk), cellobiose phosphorylase (Cbp), phosphoglucomutase (Pgmt), pyruvate phosphate dikinase (Ppdk), phosphoenolpyruvate carboxykinase (Pepck), malate dehydrogenase (Mdh), malic enzyme (Me).

Across all three glycolytic pathways, GAPDH consistently emerged as the most abundant enzyme, representing 24%, 29%, and 21% of the total glycolytic protein pool in *Z. mobilis*, *E. coli*, and *C. thermocellum*, respectively.

### 2.3 Thermodynamics shape the protein cost of glycolytic pathways

Theoretical and computational analyses predict that thermodynamically constrained reactions in glycolysis incur higher protein costs than those with larger driving forces^5,17^. These studies further suggest that glycolytic pathways with greater overall thermodynamic favorability require less protein compared to those with lower favorability. To investigate the *in vivo* relationship between pathway thermodynamics, metabolic flux, and enzyme concentration, we normalized the absolute protein concentration of each glycolytic reaction (i.e., the sum of all enzymes and isoenzymes involved) to its respective *in vivo* flux (Table S9). This approach yielded a metric of protein cost (μg protein/ (mmol hr⁻¹))^30–32^, enabling comparisons of protein costs across glycolytic reactions and pathways in the three organisms studied (Figure 3).

**Figure 3.**
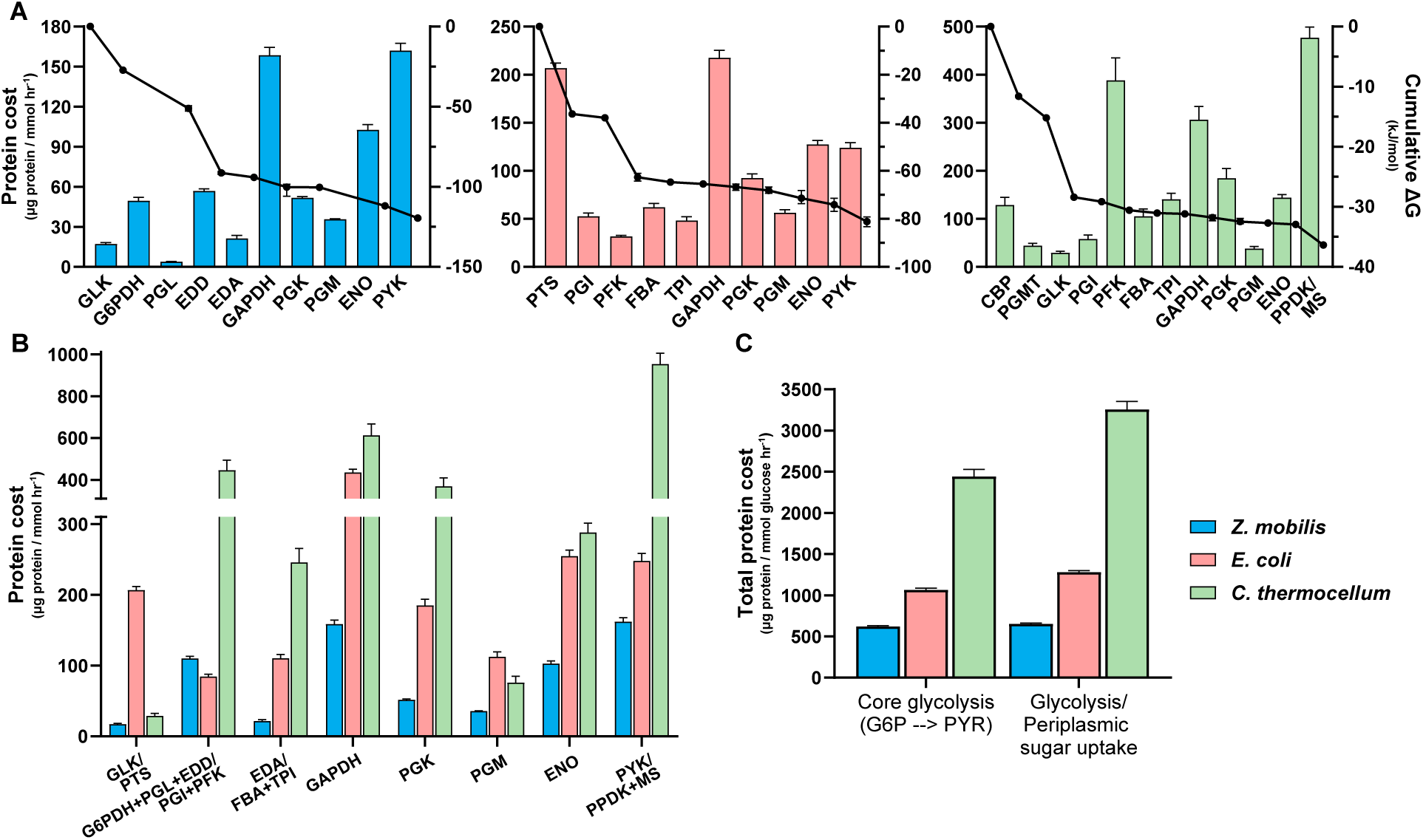
*In vivo* protein costs for glycolytic reactions and pathways. **A.** Protein costs, expressed as the amount of protein (μg) required per unit flux (mmol hr^−1^), for glycolytic reactions in *Z. mobilis* (blue), *E. coli* (pink), and *C. thermocellum* (green). The protein cost of the pyrophosphate-dependent dikinase (PPDK) and the malate shunt (MS) reactions in *C. thermocellum* were grouped together because intracellular flux measurements for the conversion of phosphoenolpyruvate (PEP) to pyruvate (PYR) do not distinguish the amount of flux occurring through each route (Table S9). The protein cost of the phosphotransferase system (PTS) in *E. coli* represents the sum of the four participating enzymes: PtsG, PtsH, PtsI, and Crr. The cumulative drop in *ΔG* (kJ mol^−1^) for each glycolytic pathway is also shown. ΔG data are a combination of previous experimental measurements^30–32^ and computationally estimated values, constrained by *in vivo* metabolite concentrations, obtained in this work (Materials and Methods). **B.** Comparison of protein costs across equivalent and analogous reactions in the ED and EMP glycolytic pathways of *Z. mobilis*, *E. coli*, and *C. thermocellum*. The aggregated protein cost for the ED reactions glucose 6-phosphate (G6P) dehydrogenase (G6PDH), phosphogluconolactonase (PGL), and 6-phosphogluconate (6PG) dehydratase (EDD) was compared to the combined protein cost for the EMP reactions phosphoglucose isomerase (PGI) and phosphofructokinase (PFK), since both sets of reactions convert G6P to the 6-carbon intermediate (either 2-keto-3-deoxy-6-phosphogluconate (KDPG) or fructose 1,6-bisphosphate (FBP)), which precedes the aldolase step. Similarly, the protein cost for the ED reaction KDPG aldolase (EDA) was compared to the combined protein cost for the EMP reactions FBP aldolase (FBA) and triose phosphate isomerase (TPI), as these include the equivalent aldolase reaction and collectively generate glyceraldehyde 3-phosphate (GAP). **C.** The total protein cost, expressed as the amount of protein (μg) required per unit flux (mmol glucose hr^−1^), for core glycolysis (G6P to PYR) and glycolysis including periplasmic sugar uptake in each bacterium. Periplasmic sugar uptake enzymes include the glucose facilitated diffusion (Glf) protein in *Z. mobilis*, the four PTS enzymes in *E. coli*, and the four enzymes (CbpB, MsdB1, MsdB2, and NbdB) that comprise transporter B in *C. thermocellum* (Table S10). Protein costs for each glycolytic reaction (panels A and B) or set of reactions (panel B) were calculated by normalizing the sum of all participating enzymes and isoenzymes to the intracellular flux of the reaction. Similarly, the protein cost for each glycolytic pathway (panel C) represents the sum of all glycolytic enzymes and isoenzymes, normalized to the glucose uptake rate of the corresponding bacterium. (Materials and Methods). For all graphs, data represent the averages of four biological replicates. Error bars show ± standard deviation (protein costs) or 95% confidence intervals (*ΔG* values). Some error bars are too small to be visible in this representation. See Table S1, S2, and S9 for *ΔG*, glucose uptake, and flux data, respectively. Abbreviations: glucokinase (GLK), glucose 6-phosphate dehydrogenase (G6PDH), phosphogluconolactonase (PGL), 6-phosphogluconate dehydratase (EDD), 2-keto-3-deoxy-6-phosphogluconate aldolase (EDA), glyceraldehyde 3-phosphate dehydrogenase (GAPDH), phosphoglycerate kinase (PGK), phosphoglycerate mutase (PGM), enolase (ENO), pyruvate kinase (PYK), phosphoglucose isomerase (PGI), phosphofructokinase (PFK), fructose 1,6-bisphosphate aldolase (FBA), triose phosphate isomerase (TPI), cellobiose phosphorylase (CBP), phosphoglucomutase (PGMT), malate shunt (MS).

Our analysis revealed a trend across all three glycolytic variants: early pathway reactions generally have lower protein costs than downstream reactions, suggesting that the initial steps operate at a higher enzyme efficiency (Figure 3A). These lower protein demands align with the larger *in vivo* thermodynamic driving forces observed in early glycolysis (Table S1). For example, in *Z. mobilis*’s ED pathway, 76% of the total change in free energy (−120 kJ mol⁻¹) occurs within the first four reactions (GLK to EDD). These reactions exhibit an average protein cost of 31.8 μg protein/(mmol hr⁻¹), nearly 3-fold lower than that of the later steps. Similarly, in the PPi-EMP pathway of *C. thermocellum*, the first three reactions (CBP to GLK) account for 80% of the pathway’s total driving force (−35 kJ mol^−1^) and have an average protein cost of 67.3 μg protein/ (mmol hr⁻¹), also about 3-fold lower than that of the subsequent reactions.

*The E. coli EMP pathway* presents a more complex scenario due to its use of the PTS, which couples glucose import to its phosphorylation to G6P while converting PEP to Pyruvate. The initial three EMP reactions (PTS, PGI, and PFK) account for 77% of the total free energy change (−83 kJ mol⁻¹). Due to the high concentrations of PTS enzymes, these reactions have an average protein cost of 97 μg protein/ (mmol hr⁻¹), comparable to the costs of lower EMP glycolytic reactions (104 μg protein/ (mmol hr⁻¹)). However, a proportion of the PTS protein cost is attributable to the conversion of PEP to pyruvate, confounding the distinction of protein costs between upper and lower glycolytic reactions. When excluding the PTS, the average protein cost of early glycolytic reactions (PGI to TPI) is about 2-fold lower than that of downstream reactions.

A central hypothesis of this study was that the higher thermodynamic favorability of the ED pathway in *Z. mobilis* would translate to lower protein costs compared to the less favorable EMP pathways in *E. coli* and *C. thermocellum*. This hypothesis was supported by our findings: lower protein costs were consistently associated with higher thermodynamic driving forces for equivalent or analogous reactions across the three glycolytic variants (Figure 3B). For the core glycolytic reactions from G6P to pyruvate, the more thermodynamically favorable ED pathway in *Z. mobilis* required approximately 4-fold and 2-fold less protein per unit flux (μg protein/ (mmol glucose hr⁻¹)) than the EMP pathways in *C. thermocellum* and *E. coli*, respectively (Figure 3C). When accounting for glucose transport systems (PTS in *E. coli* and membrane transporters in *Z. mobilis* and *C. thermocellum*), the ED pathway in *Z. mobilis* remained the most enzyme-efficient, requiring approximately 5- and 2-fold less protein per flux than the EMP pathways in *C. thermocellum* and *E. coli*, respectively (Figure 3C). These findings underscore the critical role of thermodynamic driving forces in shaping the *in vivo* protein investment required in glycolytic pathways.

### 2.4 Protein costs of sugar uptake

*Z. mobilis*, *E. coli*, and *C. thermocellum* use distinct processes for glucose or cellobiose uptake (Figure 4A). *Z. mobilis* has four carbohydrate-specific porins (OprB1, ZMO0064; OprB2, ZMO0847; OprB3, ZMOp33×009; RpfN, ZMO1859) to transport sugars across its outer membrane into the periplasm^33,54^. Among these, OprB2 was expressed at substantially higher levels compared to the other three porins (Figure 4B and Table S10). Consistent with prior studies, the glucose-facilitated diffusion protein Glf, which transports glucose from the periplasm into the cytosol, was highly expressed (Figure 4B)^55,56^. Although *Z. mobilis* encodes another transporter, ZMO0293, to import glucose into the cytosol, this protein is expressed at very low levels (Table S10), suggesting it may function under different growth conditions^57,58^.

**Figure 4.**
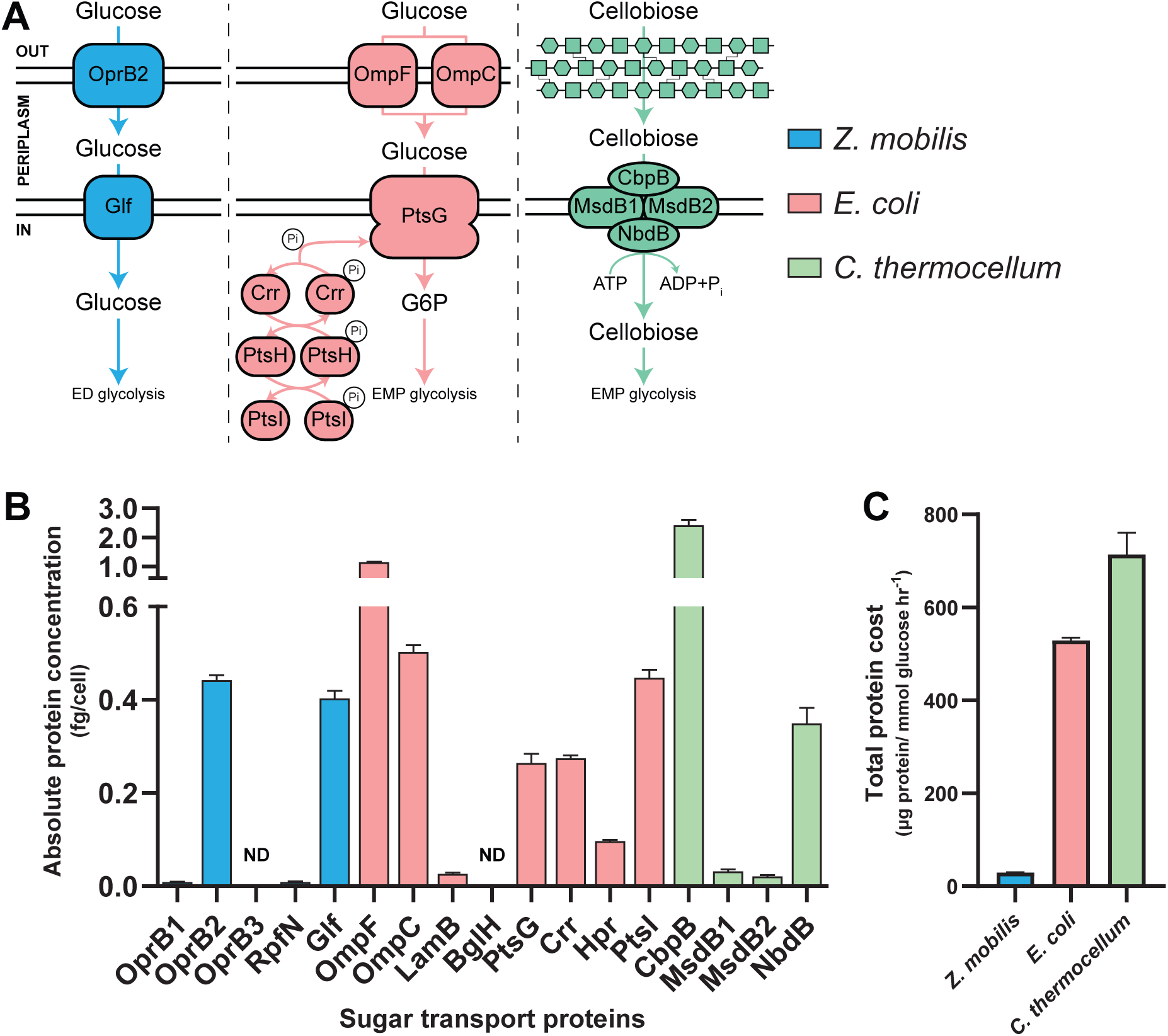
Protein costs of sugar uptake processes. **A.** *Z. mobilis* (blue), *E. coli* (pink), and *C. thermocellum* (green) use distinct enzymes and mechanisms to uptake glucose or cellobiose. In *C. thermocellum*, cellobiose enters the periplasm without the need for a dedicated transporter^115–117^. ^B^. Absolute concentrations (in fg per cell) of predominant sugar uptake proteins in each bacterium. Data are the averages of four biological replicates, with error bars indicating ± standard deviation. Enzymes designated with ND were not detected. Some error bars are too small to be visible in this representation. See Table S10 for the absolute concentration data for all sugar uptake proteins. **C.** The total protein cost of glucose uptake, expressed as the amount of protein (μg) required per unit flux (mmol glucose hr^−1^). For *C. thermocellum*, the protein cost for glucose uptake is shown as half the calculated cost for cellobiose uptake, since each molecule of cellobiose contains two glucose moieties. Abbreviations: Carbohydrate-selective porin OprB (OprB1, OprB2, OprB3), carbohydrate porin (RpfN), glucose facilitated diffusion protein (Glf), outer membrane porin F (OmpF), outer membrane porin C (OmpC), Maltoporin (LamB), Cryptic outer membrane porin (BglH), Phosphotransferase system (PTS) glucose-specific EIICB component (PtsG), PTS system glucose-specific EIIA component (Crr), Phosphocarrier protein HPr (PtsH), Phosphoenolpyruvate-protein phosphotransferase (PtsI), extracellular solute-binding protein family 1 (CbpB), binding-protein-dependent transport systems inner membrane component (MsdB1, MsdB2), ABC transporter related protein (NbdB).

*E. coli* has four outer membrane porins (OmpF, b0929; OmpC, b2215; BglH, b3720; LamB, b4036) to transport sugars into its periplasm^59^. Consistent with previous studies showing that OmpF and OmpC are utilized for glucose uptake, these two porins were highly expressed relative to LamB and BglH (Figure 4B and Table S10)^60–62^. Periplasmic glucose is subsequently transported into the cytosol via the PTS, which consists of four phospho-relay proteins: PtsG, b1101; Hpr, b2415; PtsI, b2416; Crr, b2417^63–65^. Our data confirm that all four PTS components are highly expressed under the conditions tested (Figure 4B).

*C. thermocellum* harbors five multi-component ATP-binding cassette (ABC) transporters to import sugars across the cell membrane: transporters A, B, C, D, and L^66^. Consistent with previous research showing that *C. thermocellum* primarily uses transporter B to uptake cellobiose, our data show that the components of transporter B (MsdB1, Clo1313_1195; MsdB2, Clo1313_1196; NbdB, Clo1313_2554; CbpB, Clo1313_1194) are expressed at much higher levels than the other ABC transporter proteins (Table S10)^67^. Notably, the cellobiose binding protein CbpB (Clo1313_1194) was the second most abundant protein in the *C. thermocellum* proteome, while the transmembrane (MsdB1/2) and ATP binding (NbdB) subunits were expressed at lower levels (Figure 4B and Table S8).

Our analysis reveals that *E. coli* and *C. thermocellum* allocate significantly more protein (∼3 fg cell^−1^ each) for glucose or cellobiose uptake than *Z. mobilis* (∼1 fg cell^−1^). Furthermore, due to its higher glucose uptake rate (Table S2), *Z. mobilis* requires more than 10 times less protein to import an equivalent amount of glucose compared to *E. coli* and *C. thermocellum* (Figure 4C).

### 2.5 Protein burden of fermentative pathways is influenced by reversibility

*Z. mobilis* and *C. thermocellum* produce ethanol and acetate via distinct metabolic routes (Figure 5A). In *Z. mobilis*, over 95% of carbon is directed towards ethanol, with minimal production of other fermentation products, such as acetate, formate, and lactate^68,69^. Reflecting this, the concentrations of the ethanol fermentation enzymes pyruvate decarboxylase (Pdc) and alcohol dehydrogenase B (AdhB) are much higher compared to those involved in acetate or lactate production (Figure 5B and Table S11)^70,71^. Notably, the levels of Pdc and AdhB are comparable to those of glycolytic enzymes, with Pdc levels exceeding those of all glycolytic enzymes on a mass basis (fg cell^−1^) (Table S4).

**Figure 5.**
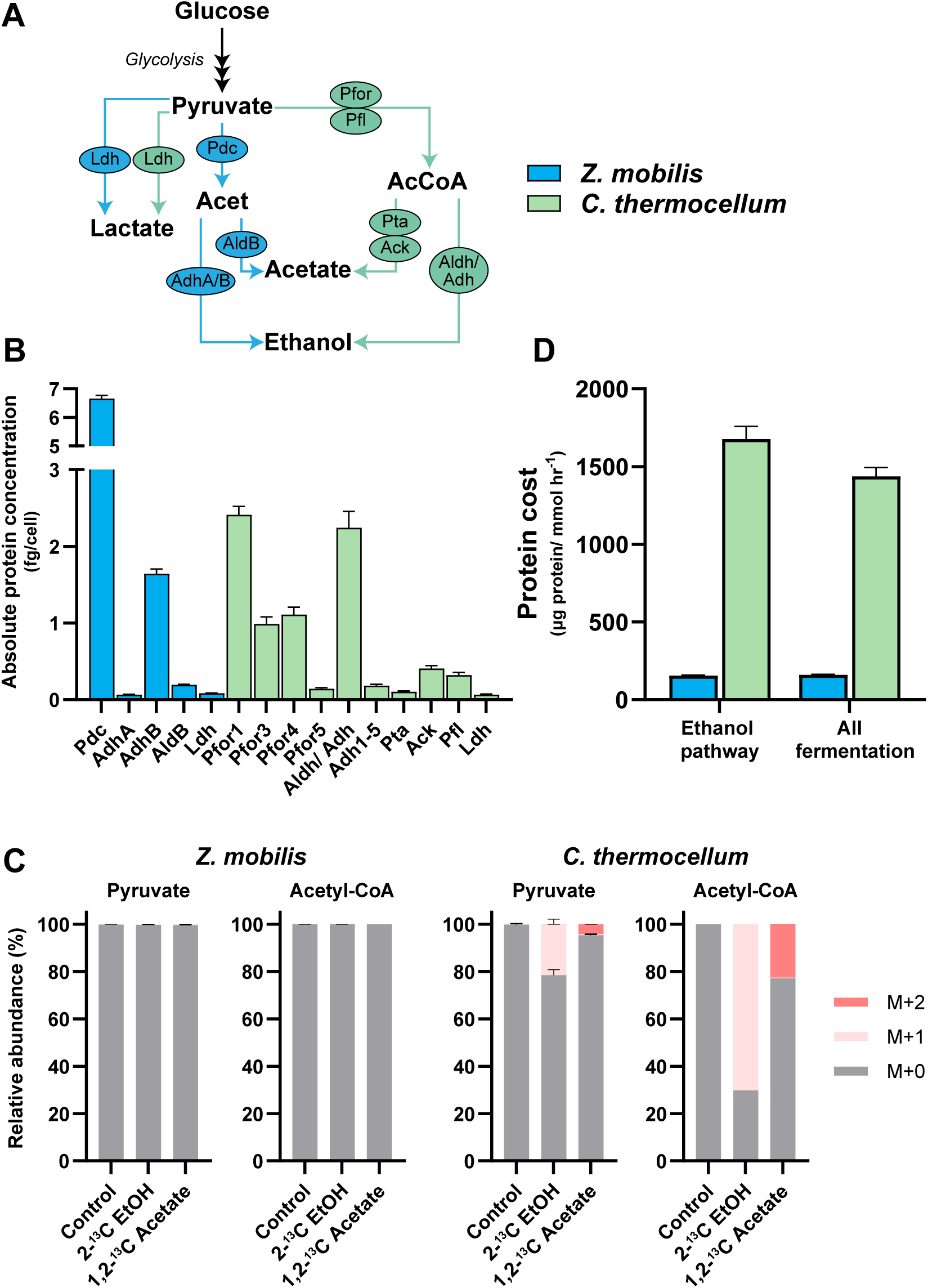
Protein costs and reversibility of fermentation pathways in *Z. mobilis* and *C. thermocellum*. **A.** Fermentation pathways in *Z. mobilis* (blue) and *C. thermocellum* (green). **B.** Absolute concentrations of fermentation enzymes and complexes, expressed in fg per cell. Data represent individual protein concentrations (e.g., Pdc, AdhA, AdhB, etc.), the sum of protein subunits forming a complex (e.g., Pfor1, PFor4, etc.), the combined protein and its activator (i.e., Pfl), or the sum of all enzymes and isoenzymes participating in the reaction (e.g., Ldh, Adh1-5) (see Table S11). **C.** Labeling patterns of pyruvate and acetyl-CoA when cells are grown in the presence of isotopically labeled ethanol (EtOH) or acetate. The *y*-axis represents the fraction of the metabolite pool that is isotopically labeled, corrected for the natural abundance of ^13^C (Materials and Methods). M+0 indicates all atoms are ^12^C, and M+1 indicates one ^13^C atom is present. Pyruvate labeling patterns were calculated from valine to exclude unlabeled (M+0) pyruvate originating from the media. Acetyl-CoA labeling specifically refers to the acetyl group, calculated from aspartate and glutamate labeling data. **D.** Protein costs of fermentation, expressed as the amount of protein (μg) required per unit flux (mmol glucose hr^−1^) in each bacterium. Protein costs were calculated by normalizing the sum of ethanol fermentation enzyme concentrations to ethanol flux (left bars, Ethanol pathway), or by normalizing the sum of ethanol, acetate, formate, and lactate fermentation enzyme concentrations to total fermentation flux (bars on the right, Total fermentation) (Materials and Methods). Data represent the averages of four biological replicates. Error bars show ± standard deviation. Some error bars are too small to be visible in this representation. See Table S11 for absolute concentration data for fermentation and acetyl-CoA metabolism proteins. Abbreviations: lactate dehydrogenase (Ldh), pyruvate decarboxylase (Pdc), pyruvate ferredoxin oxidoreductase (Pfor), pyruvate formate-lyase (Pfl), alcohol dehydrogenase (Adh, AdhA/B), NADP^+^-dependent acetaldehyde dehydrogenase (AldB), phosphate acetyltransferase (Pta), acetate kinase (Ack), bifunctional acetaldehyde/ alcohol dehydrogenase (Aldh/ Adh).

*C. thermocellum* produces ethanol and acetate as its primary fermentation products^72^. Ethanol fermentation in *C. thermocellum* involves pyruvate ferredoxin oxidoreductase (Pfor), pyruvate formate-lyase (Pfl), and the bifunctional acetaldehyde/ alcohol dehydrogenase (Aldh/ Adh) enzymes (Figure 5A)^73,74^. While *C. thermocellum* encodes five annotated Pfor complexes (Table S11), previous studies have shown that deletion of Pfor1 (Clo1313_0020-0023) or Pfor4 (Clo1313_1353-1356) reduces PFOR activity by 80%, suggesting these two complexes play a major role in ethanol fermentation ^75,76^. Consistent with these findings, we observed that the subunits of Pfor1 and Pfor4 are the most abundant among all the Pfor complexes, but we also found that Pfor3 subunits are highly expressed as well (Figure 5B). Notably, the abundances of these Pfor complexes and Aldh/ Adh are comparable to that of highly abundant glycolytic enzymes (Table S8) and exceeds the concentrations of enzymes involved in acetate and lactate production (Figure 5B and Table S11).

Previous studies suggest that the PFOR reaction in *C. thermocellum* is highly reversible^77–80^. In contrast, Pdc in *Z. mobilis* catalyzes a reaction that is considered to have limited reversibility^30,81^. We hypothesized that the ethanol and acetate fermentation pathways in *C. thermocellum*, which are preceded by a glycolytic pathway with limited thermodynamic driving force, are highly reversible and thermodynamically constrained. Conversely, the ethanol fermentation pathway in *Z. mobilis*, reliant on the PDC reaction and preceded by the thermodynamically favorable ED pathway, is expected to be largely irreversible. While previous computational thermodynamic analyses have supported these hypotheses^80^, direct experimental evidence regarding the reversibility of fermentation pathways in *C. thermocellum* and *Z. mobilis* is lacking.

To investigate the reversibility of these fermentation pathways, we cultured *Z. mobilis* and *C. thermocellum* in the presence of 2-^13^C-labeled ethanol and 1,2-^13^C-labeled acetate (Materials and Methods) and tracked the incorporation of isotope labeling into upstream metabolites, including pyruvate and acetyl-CoA. In *Z. mobilis*, there was no detectable incorporation of ^13^C from labeled ethanol or acetate into acetyl-CoA or pyruvate, indicating that its ethanol and acetate fermentation pathways are highly irreversible (Figure 5C). In contrast, *C. thermocellum* showed substantial incorporation of ^13^C from labeled ethanol and acetate into acetyl-CoA, with lesser incorporation into pyruvate (Figure 5C). Specifically, 70% of acetyl-CoA was labeled from ^13^C-ethanol, reflecting the high reversibility of the ALDH/ ADH reactions, while 20% of pyruvate was labeled, highlighting the reversibility of PFOR/ PFL reactions (Figure 5C). Similarly, when *C. thermocellum* was cultured on ¹³C-labeled acetate, we observed significant labeling of acetyl-CoA (23%) and a smaller fraction of labeled pyruvate (5%), indicating that the phosphate acetyltransferase (PTA) and acetate kinase (ACK) reactions are also reversible (Figure 5C), and further supporting the reversibility of PFOR/PFL. These findings demonstrate that, similar to its glycolytic pathway, the ethanol and acetate fermentation pathways in *C. thermocellum* are highly reversible and thermodynamically constrained.

Given the pronounced differences in the reversibility of ethanol and acetate fermentation pathways between *Z. mobilis* and *C. thermocellum*, we predicted that the protein cost associated with these pathways would be substantially higher in *C. thermocellum*. Supporting this hypothesis, our analysis showed that the total protein cost for fermentation reactions was 9-fold higher in *C. thermocellum* than in *Z. mobilis* to ferment an equivalent amount of glucose into ethanol and acetate (Figure 5D). For ethanol production alone, *C. thermocellum* required nearly 11-fold more protein compared to *Z. mobilis*. Thus, similar to its glycolytic pathway, *C. thermocellum* incurs a higher enzymatic cost for fermentation compared to *Z. mobilis* due to the limited thermodynamic driving force of its pathways.

### 2.6 Proteome-wide allocation of protein resources

To examine how the bacteria studied allocate their protein resources across cellular processes, we conducted a Cluster of Orthologous Groups (COG) analysis (Materials and Methods), classifying proteins into distinct biological functions (e.g., transcription, cell motility, carbohydrate metabolism and transport, etc.) (Tables S6-S8, S12) and quantifying protein allocation to each category^82,83^. Overall, protein allocation across COG-defined cellular functions was largely consistent across the organisms studied. Five major categories −translation, ribosomal structure and biogenesis; amino acid transport and metabolism; energy production and conversion; carbohydrate transport and metabolism; and cell wall/membrane/envelope biogenesis− accounted for 58%, 70%, and 59% of the proteome (on a fg cell^−1^ basis) in *Z. mobilis*, *E. coli*, and *C. thermocellum*, respectively (Figure 6).

**Figure 6.**
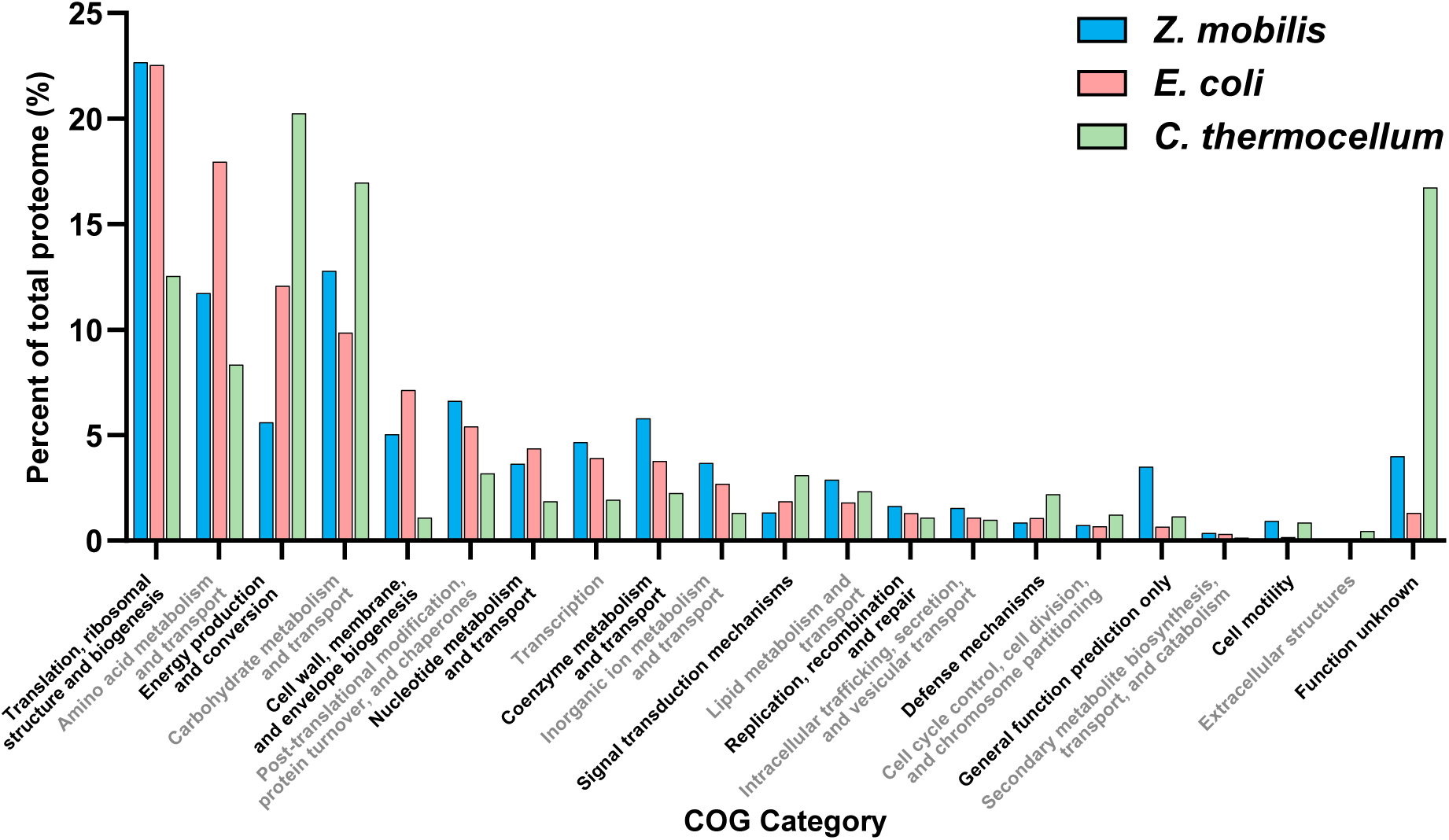
Allocation of protein resources to cellular processes. Proportion of the measured proteome dedicated to 21 COG-defined cellular processes in *Z. mobilis* (blue), *E. coli* (pink), and *C. thermocellum* (green). COG categories accounting for less than 0.1% of total protein mass are excluded. Protein mass was quantified in femtograms per cell (fg cell⁻¹). COG classifications for each protein are provided in Tables S6-S8, and the corresponding letter designations for COG categories can be found in Table S12.

Despite these broad similarities, we also observed notable differences. In *Z. mobilis* and *E. coli,* the cellular process with the largest allocation of protein resources was protein biogenesis (i.e., translation, ribosomal structure/ biogenesis), comprising ∼23% of the proteome in both bacteria. In contrast, the most resource-demanding process in *C. thermocellum* was energy production and conversion, accounting for 20% of its proteome. This was followed closely by carbohydrate metabolism and transport, which constituted 17% of the proteome (Figure 6).

## 3. Discussion

This study provides *in vivo* evidence that thermodynamic driving forces are a key determinant of enzyme burden in metabolic pathways. We show that the more thermodynamically favorable ED glycolytic pathway in *Z. mobilis* requires substantially less enzymatic protein to sustain the same flux as the less favorable PPi-EMP and EMP pathways in *C. thermocellum* and *E. coli*. Additionally, we show that the highly reversible fermentation pathways in *C. thermocellum* impose a markedly higher protein cost compared to the irreversible fermentation pathways in *Z. mobilis*.

### Comparison with theoretical predictions and trade-offs between ATP yield and protein cost

A previous theoretical analysis, based on computationally-derived thermodynamic values and *in vitro* enzyme kinetics data, estimated that the canonical EMP pathway requires between 3.5-to 5-fold more enzymatic protein than the ED pathway to sustain the same glycolytic flux^17^. Our experimental findings closely align with these predictions, as we found that the ED pathway in *Z. mobilis* requires approximately 5- and 2-fold less protein to achieve the same glycolytic flux as the EMP pathways in *C. thermocellum* and *E. coli*, respectively (Figure 3C).

The ED pathway in *Z. mobilis* generates 1 ATP per glucose, while the EMP pathway in *E. coli* produces 2 ATPs, and the PPi-EMP pathway in *C. thermocellum* yields 4 ATP equivalents^22,84^. Our findings provide strong experimental support for the predicted tradeoff between glycolytic ATP yield and protein costs. The higher energy yield of the PPi-EMP pathway in *C. thermocellum*, linked to its limited thermodynamic driving force, comes at the expense of significantly greater enzyme burden relative to the EMP and ED pathways in *E. coli* and *Z. mobilis*. This reliance on a thermodynamically constrained glycolytic pathway with increased ATP yield is likely an evolutionary adaptation to growth on cellulosic substrates. Microorganisms metabolizing soluble substrates can optimize either a high specific substrate consumption rate (grams of substrate consumed per gram of cells per hour) or a high cell yield (grams of cells produced per gram of substrate), both of which contribute to maximizing the specific growth rate. However, for microbes growing on cellulosic biomass, the specific substrate consumption rate is inherently limited, creating strong selective pressure to maximize cell yield by increasing glycolytic ATP yield^31^. In contrast, highly thermodynamically favorable pathways with lower ATP yield, such as the ED pathway in *Z. mobilis*, are well-suited for environments rich in glucose, where rapid substrate consumption provides a competitive advantage.

### High protein cost of PTS in *E. coli*

Across the glycolytic pathways examined, we observed a general trend in which early reactions with greater thermodynamic favorability incur lower protein costs, whereas less favorable downstream steps require greater enzyme investment. However, we identified several exceptions to this trend. One such exception is the PTS in *E. coli*, which, despite being highly thermodynamically favorable (Table S1), incurs one of the highest protein costs within EMP glycolysis (Figure 3A). This elevated cost likely arises from its dual role in both catalysis and regulation. Specifically, the PTS is involved in carbon catabolite repression and inducer exclusion, regulating the uptake of preferred carbon sources^85^. Additionally, only a fraction of catalytic components may be active at any given time due to feedback inhibition. For example, the *E. coli* PTS is inhibited by α-ketoglutarate, a TCA cycle intermediate involved in nitrogen assimilation that binds non-competitively to PtsI preventing PEP dephosphorylation^86^. Notably, PtsI in the most highly expressed PTS component (Table S10). These regulatory roles likely necessitate abundant expression of PTS components, thereby increasing its overall protein cost.

Another notable exception is the PGM reaction in *C. thermocellum*, which, despite operating near equilibrium (≤ −0.48 kJ mol^−1^)^31^, exhibits one of the lowest protein costs within the pathway (Figure 3A). *C. thermocellum* encodes multiple Pgm isoenzymes (Pgm1-Pgm7), with Pgm1 being the most highly expressed, but still maintains a low overall protein burden for this reaction. (Tables S3 and S4). This could be explained by high catalytic efficiency or high active site saturation, but direct measurements are unavailable. Alternatively, an unannotated, highly expressed enzyme might be responsible for catalyzing this step.

### Metabolic engineering strategies to reduce protein costs in *C. thermocellum*

The use of a PPi-dependent phosphofructokinase (PPi-Pfk) in *C. thermocellum*’s glycolytic pathway enables the generation of one additional ATP per glucose compared to the ATP-dependent phosphofructokinase (ATP-Pfk) employed in the EMP pathway of *E. coli*. However, this increased ATP yield comes with a significant tradeoff: the protein cost of the PFK reaction in *C. thermocellum* is more than 12-fold higher than in *E. coli*.

Given its exceptional ability to degrade cellulose, *C. thermocellum* is widely regarded as a promising platform organism for consolidated bioprocessing of lignocellulosic biomass into fuels and chemicals^87^. One potential strategy to improve biofuel production in this organism involves reducing the high protein cost of the PPi-EMP pathway by alleviating thermodynamic bottlenecks through metabolic engineering. A recent study demonstrated the feasibility of this approach by replacing PPi-Pfk with ATP-Pfk, deleting *ppdk*, and introducing genes encoding a soluble pyrophosphatase (PPase) and pyruvate kinase (Pyk) to engineer a PPi-free glycolytic pathway in *C. thermocellum*^88^. These modifications improved the thermodynamics of the PFK reaction and increased ethanol titers by 38%. Future studies could assess how these modifications impact enzyme efficiency by quantifying protein levels in the engineered pathway. Furthermore, given the high protein cost and limited thermodynamic driving force of the ethanol fermentation pathway in *C. thermocellum*, another potential strategy to enhance ethanol production could involve increasing the thermodynamic favorability of this pathway, potentially by replacing Pfor with pyruvate decarboxylase (Pdc).

### Enhanced glycolytic flux during N₂ fixation in *Z. mobilis*

A potential drawback of the highly forward driven ED pathway in *Z. mobilis* is that when cellular demand for energy or biomass increases, each of the enzyme-efficient steps in the pathway can become a kinetic bottleneck, potentially requiring an increase in enzyme concentration to increase flux. Interestingly, *Z. mobilis* can increase its glycolytic rate by approximately 40%, compared to growth under ammonia-replete conditions, when utilizing dinitrogen gas (N_2_) as its sole nitrogen source^89–91^. This increase in glycolytic rate correlates with increased thermodynamic favorability of the ED pathway^92^.

Leveraging previous proteomics data^92^, we compared ED pathway enzyme levels and protein costs between N₂-fixing and NH₄⁺-replete conditions (Figure S2, Table S13). While the levels of most ED pathway enzymes remain unchanged or increased only marginally during N₂-fixing conditions, Pgl, Pgk, and Eno displayed significant increases (Figure S2A). Notably, Pgl, the least abundant ED enzyme under NH₄⁺-replete conditions, showed a 1.8-fold increase during N_2_ fixation, suggesting that this enzyme may be a rate-limiting step. Normalizing enzyme levels to intracellular fluxes revealed that most ED enzymes —including all enzymes in the lower half of the pathway— exhibited significantly lower protein costs under N₂-fixing conditions (Figure S2B), indicating that *Z. mobilis* utilizes its glycolytic enzymes more efficiently when nitrogen availability is limited.

The increased thermodynamic driving force of lower ED pathway reactions under N₂-fixing conditions likely contributes to decreased protein costs and higher flux for these enzymes; in contrast, regulatory mechanisms −such as allosteric control or post-translational modifications− might be responsible for improving enzyme efficiency and flux of the early highly thermodynamically favorable steps of the pathway^93–95^. For example, the *ΔG* of GAPDH improves from −0.90 to −1.62 kJ mol^−1^ during N_2_ fixing conditions^92^. Despite this seemingly minor increase in thermodynamic favorability, this change in free energy corresponds to a 1.8-fold higher net flux (Table S15). Similarly, the *ΔG* of EDA decreases by 0.46 kJ mol^−1^ (−1.42 to −1.88 kJ mol^−1^)^92^, which corresponds to a 1.3-fold higher net flux (Table S15). These observations align with a previous study in *E. coli* showing that increases in glycolytic rates during nitrogen or phosphorus upshift correlate with increased thermodynamic driving force of pathway steps initially close to equilibrium^96^.

However, the highly thermodynamically favorable upper ED pathway reactions (i.e., GLK, G6PDH, PGL, and EDD) already operate with very high efficiency, so increases in thermodynamic favorability are predicted to have a negligible effect on net flux. For the PGL reaction, increased flux could be explained by the 1.8-fold increase in Pgl levels, but for the other upper ED pathway reactions, whose levels don’t increase significantly, other explanations are warranted. Considering thermodynamically favorable reactions in glycolysis are known to be targets of metabolic regulation in several organisms^97^, another possibility as to how *Z. mobilis* cells are capable of sustaining enhanced flux through glycolytic reactions under N_2_ fixing conditions is by modulating enzyme activity via metabolic regulation and/ or post-translational modifications^94,95^. Notably, phospho-proteomic analyses have identified phosphorylation changes in multiple *Z. mobilis* glycolytic enzymes under N₂-fixing conditions^93^. Additional research is needed to determine if these post translational modifications regulate enzyme activity and contribute to higher glycolytic flux.

### Conclusion

This study provides *in vivo* evidence that thermodynamic driving forces play a major role in shaping enzyme burden in glycolytic pathways. The insights and quantitative proteomic data generated here will serve as a valuable resource for developing constraint-based genome-scale metabolic models, such as resource balance analysis (RBA) and metabolism and expression (ME) models, which explicitly account for the amount of enzyme needed to sustain metabolic flux.

## 4. Materials and Methods

### Strains and growth conditions

#### Zymomonas mobilis

ZM4 (ATCC 31821) was streaked onto *Zymomonas* rich-medium glucose (ZRMG) plates (10 g/L yeast extract, 2 g/L KH_2_PO_4_, 20 g/L glucose, and 20 g/L agar) from 25% glycerol stocks and incubated in an anaerobic (5% H_2_, 5% CO_2_, 90% N_2_ atmosphere, and <100 ppm O_2_) chamber (Coy Laboratory) at 30°C for 3 to 4 days. Single colonies were used to inoculate 15 mL test tubes containing 10 mL liquid ZRMG. Cells were grown overnight and then subcultured into 25-50 mL of *Zymomonas* minimal media (ZMM) [1 g/L K_2_HPO_4_, 1 g/L KH_2_PO_4_, 0.5 g/L NaCl, 1 g/L (NH_4_)_2_SO_4_, 0.2 g/L MgSO_4_·6H_2_O, 0.025 g/L Na_2_MoO_4_·2H_2_O, 0.0025 g/L FeSO_4_·7H_2_O, 0.02 g/L CaCl_2_·2H_2_O, 5 mg/L calcium pantothenate, and 20 g/L glucose]. These subcultures were incubated for 14-16 hours and used to inoculate experimental cultures.

#### Escherichia coli

RL3000 (MG1655 *ilvG^+^ rph^+^ pyrE^+^*), a non-hyperflagellated prototrophic derivative of MG1655^98^ was streaked onto Luria-Broth (LB) plates (10 g/L tryptone, 5 g/L yeast extract, 5 g/L NaCl, 15 g/L agar) from 25% glycerol stocks and incubated aerobically at 37 °C for 16-18 hours. Single colonies were used to inoculate 15 mL test tubes containing 10 mL of liquid LB. Cells were grown for 8-10 hours at 37 °C 250 RPM and then subcultured into 25-50 mL of M9 minimal media (6 g/L Na_2_HPO_4_, 3 g/L KH_2_PO_4_, 0.5 g/L NaCl, 1 g/L NH_4_Cl, 0.12 g/L MgSO_4_, 0.0147 g/L CaCl_2_, 0.002 g/L FeSO_4_·7H_2_O, and 4 g/L glucose). These subcultures were incubated for 14-16 hours and used to inoculate experimental cultures.

#### Clostridium thermocellum

DSM1313 growth was carried out anaerobically in MTC media (9.39 g/L morpholine propanesulfonic acid [MOPS] sodium salt, 2 g/L potassium citrate monohydrate, 1.3 g/L citric acid monohydrate, 1 g/L Na_2_SO_4_, 1 g/L KH_2_PO_4_, 2.5 g/L NaHCO_3_, 2 g/L urea, 1 g/L MgCl_2_·6H_2_O, 0.2 g/L CaCl_2_·2H_2_O, 0.1 g/L FeCl_2_·4H_2_O, 1 g/L L-cysteine HCl monohydrate, 0.02 g/L pyridoxamine HCl, 0.004 g/L *p*-aminobenzoic acid [PABA], 0.002 g/L biotin, 0.002 g/L vitamin B_12_, 0.004 g/L thiamine, 0.5 µg/L MnCl_2_·4H_2_O, 0.5 µg/L CoCl_2_·6H_2_O, 0.2 µg/L ZnCl_2_, 0.1 µg/L CuCl_2_·2H_2_O, 0.1 µg/L H_3_BO_3_, 0.1 µg/L Na_2_MoO_4_·2H_2_O, 0.1 µg/L NiCl_2_·6H_2_O, and 5 g/L cellobiose). To prepare MTC media, tubes or bottles were filled with an initial base media containing MOPS solution, sealed with butyl rubber stoppers, made anaerobic via a vacuum manifold, overlaid with N_2_ gas (oxygen scrubbed), and autoclaved. The additional media components were made anaerobic, autoclaved separately, and then added to the culture tubes/ bottles. Before inoculating/ extracting cultures, syringes were made anoxic by multiple drawings and expulsions of the headspace from an anaerobic sealed bottle containing 2.5% cysteine HCl solution. Cultures were inoculated directly from 25% glycerol stocks into 5 mL of MTC media and grown anaerobically in a 55 °C water bath for 24 hrs. Cultures were then subcultured into 10 mL of fresh MTC and grown for 14-16 hours, and the subcultured growth was used to inoculate experimental cultures. Experimental cultures for all three microbes were inoculated at an initial OD_600_ of 0.05 to 0.06.

### Protein extraction and sample preparation for proteomics analyses

When cells reached an OD_600_ of 0.45-0.46, 10 mL of bacterial culture was collected in a pre-chilled 15 mL conical tube for four biological replicates and centrifuged at 4,255 × g for 5 mins at 4 °C. Cell pellets were washed with phosphate buffered saline (8 g/L NaCl, 0.2 g/L KCl, 1.44 g/L Na_2_HPO_4_, 0.24 g/L KH_2_PO_4_)^99^, centrifuged at 16,000 × *g* for 5 min at 4°C, and the supernatant was discarded. Cell pellets were then stored at −80 °C until proteomics analysis.

Cell pellets were thawed and resuspended in denaturing buffer (5.4 M guanidinium hydrochloride, 100 mM Tris HCl). Samples were sonicated for 5 minutes in a chilled water bath (QSonica) using the following program: 20 seconds on, 10 seconds off, amplitude of 30, and temperature maintained at 14 °C. Samples were then incubated in a sand bath at 110 °C for 5 minutes, cooled at room temperature for 5 minutes, and incubated again in the sand bath at 110°C for 5 minutes. To precipitate the protein, liquid chromatography mass spectrometry (LC-MS) grade MeOH was added to each sample to a final solution volume of 90% MeOH v/v and vortexed. Samples were centrifuged at 14,000 x g for 2 minutes at 4 °C to pellet precipitated protein, and the supernatant was carefully removed without disturbing the protein pellet. Protein pellets were resuspended in 8 M urea, 100 mM Tris HCl, 10 mM TCEP, 40 mM chloroacetamide and vortexed for 10 minutes at room temperature to resolubilize the protein. In a 1:50 protease:protein mass ratio, 1 mg/mL LysC prepared per manufacturer’s instruction (VWR, Radnor, PA) was added to each sample, then incubated at ambient for four hours with gentle rocking. Samples were then diluted with freshly prepared 100 mM Tris HCl, pH 8.0 in order to bring the sample urea concentration to 2 M. Trypsin (Promega, Madison, WI) was added to each sample at a 1:50 protease:protein mass ratio, and samples were incubated at ambient temperature overnight while gently rocking. The digestion reaction was terminated by adding sufficient 10% TFA in H_2_O to each solution to bring solution pH to <2, as verified by pH strip.

Samples were centrifuged at 14,000 x g for 2 minutes at ambient to pellet insoluble material. Resulting supernatant was desalted using Strata-X 33 µm polymeric reversed phase SPE cartridges (Phenomonex, Torrance, CA). The desalted peptides were dried down in a vacuum centrifuge (Thermo Fisher Scientific, Waltham, MA). Peptides were resuspended in water to determine peptide concentration via NanoDrop One Microvolume UV-Vis spectrophotometer (Thermo Fisher Scientific, Waltham, MA). For samples used for absolute protein quantification, peptides were combined with synthetic HeavyPeptide AQUA peptide standards (Thermo Fisher Scientific, Rockford, IL). For each sample, two dilutions were prepared to ensure peptide standard concentrations were approximately close to the native peptide concentrations as estimated by shotgun proteomic analyses. The sample mixtures were dried down again, then resuspended in 40% acetonitrile in 0.2% formic acid for infusion. For samples used for LC-MS shotgun proteomics analysis, desalted peptides were resuspended in 0.2% formic acid and peptide concentrations were quantified via NanoDrop.

### Absolute proteomics MS and shotgun proteomics LC-MS methods

Sample analysis for absolute protein quantitation was performed using the TriVersa NanoMate (Advion, Ithaca, NY) coupled to an Orbitrap Eclipse Tribrid mass spectrometer (Thermo Fisher Scientific, San Jose, CA). The NanoMate was equipped with a 5 µm nominal internal diameter nozzle ESI chip operated at 1.60 kV, with a gas pressure of 1.0 psi, and 10 µL injection volume, with remaining volume returned to well after an injection. The MS was operated in positive ionization mode via parallel reaction monitoring (PRM), in which the m/z values corresponding to the ions from the native and isotope-labelled peptides were targeted for MS2 spectral acquisition. Targeted precursor ions were isolated from a 0.5 Da isolation window in the quadrupole; HCD MS2 scans with 25% fixed collision energy and a normalized AGC target (%) of 200, equivalent to 1e5 ions, were collected in the Orbitrap from 350-2,000 m/z with a resolution of 500,000. Maximum injection time was set to 1,014 ms for higher concentration samples or 5,000 ms for lower concentration samples.

To analyze samples for shotgun LC-MS proteomics, 2 μg of peptides was loaded onto a 75-μm-inside-diameter (i.d.), 30-cm-long capillary with an imbedded electrospray emitter and packed in a 1.7-μm-particle-size C_18_ BEH column. The mobile phases used were as follows: phase A, 0.2% formic acid; and phase B, 0.2% formic acid–70% acetonitrile. Peptides were eluted with a gradient increasing from 0% to 75% B over 42 min followed by a 4-min 100% B wash and 10 min of equilibration in 100% A for a complete gradient of 60 min.

The eluting peptides were analyzed with an Orbitrap Fusion Lumos (Thermo Scientific) mass spectrometer. Survey scans were performed at a resolution of 240,000 with an isolation analysis at 300 to 1,350 *m/z* and AGC target of 1e6. Data-dependent top-speed (1-s) tandem MS/MS sampling of peptide precursors was enabled with dynamic exclusion set to 10 s on precursors with charge states 2 to 4. MS/MS sampling was performed with 0.7-Da quadrupole isolation and fragmentation by higher-energy collisional dissociation (HCD) with a collisional energy value of 25%. The mass analysis was performed in the ion trap using the “turbo” scan speed for a mass range of 200 to 1,200 *m/z*. The maximum injection time was set to 11 ms, and the AGC target was set to 20,000.

### Absolute and shotgun proteomics data analysis

For targeted data analysis, raw data files from the PRM direct infusion-MS/MS experiments were imported into Skyline 22.2.0.351. Three to five transitions per targeted precursor ion were manually integrated to quantitate over a period of time where the MS2 ion intensities were stable. For a given native peptide and its matching isotope-labelled peptide, selected transitions were quantitated over the same period of time.

Data were exported into Excel. For each quantitated transition, the measured area was divided by the length of time over the quantitation to calculate the height. The calculated height values were summed for each set of transitions per precursor ion. If multiple charge states were tracked for a peptide (e.g. 2+ and 3+), these summed height values were added together. These summed values, as well as the known concentration of the isotope-labeled peptide spiked into the sample mixture, were consequently used to calculate the concentration of the native peptide, with corrections for dilution as necessary. Concentration data were normalized to a per cell or mass basis using calculated cell numbers, volumes, and grams per dry cell weight (gDCW) measurements (Table S2). Normalized root mean square errors (NRMSE) across peptides and biological replicates were calculated using the equation: 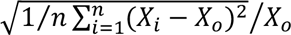 (Table S4).

Raw shotgun LC-MS proteomics data were analyzed using the MaxQuant software (version v2.6.2.0)^100^. Spectra were searched using the Andromeda search engine against a target decoy database. FASTA reference proteomics for each microbe were obtained from the National Center for Biotechnology Information (NCBI) or UniProt databases. Label free quantitation (i.e., iBAQ) was toggled on, and default values were used for all other analysis parameters. The peptides were grouped into subsumable protein groups and filtered to reach 1% false discovery rate (FDR) based on the target decoy approach. iBAQ intensities were log_2_-transformed, and these values for proteins that were absolutely quantified were used to construct a quantitative model for global protein quantification (Figure S1). Cross validation of our quantitative model was performed using the leave one out cross validation method via the R package caret (version 6.0.94).

Absolute protein concentrations for *Z. mobilis* grown under N_2_ fixation conditions were quantified using label-free quantification (LFQ) proteomics data from a previous study^92^. Growth conditions for NH_4_ replete *Z. mobilis* cells grown in Martien *et al* 2021 were identical to the growth conditions used in our absolute proteomic measurements^92^. Thus, we normalized the NH_4_ replete proteomics data from Martien *et al* 2021 to absolute values and used the fold-change measurements between NH_4_ replete and N_2_ fixation conditions to obtain absolute values for *Z. mobilis* proteins when grown under N_2_ fixation conditions (Table S13)^92^.

### Cell volume measurements via microscopy

To calculate cell volumes, 1 mL of cells was collected in a tube at an OD_600_ of 0.45-0.46 following the previously described growth scheme. The tube containing cells was not placed on ice to minimize fluctuations in cell volume induced by temperature changes. Following sample collection, 1 µL of cells was placed on a 1.5% agarose pad made with M9, ZMM, or MTC media without carbon source to reduce cell movement, and the pad was placed cell-side down onto a coverslip. Samples were analyzed < 5 minutes following removal from the flask via phase-contrast microscopy with a resolution of 0.1083 µm/pixel using a 60X UPlanSApo oil objective attached to an Olympus IX83 inverted microscope and an ORCA-Flash4.0 V2 digital camera (Hamamatsu, C11440-22CU). Cell images were adjusted for brightness and contrast and analyzed using the ImageJ software^101^. For each bacterium, length (L) and width (W) dimensions of 100 individual cells were obtained, and cell volumes (V) were calculated by assuming cells have the shape of a cylinder capped with two half-spheres and the following formula: *V* = (*π* · *W*^2^) · (*L* − *W*/3)⁄4 (Table S2)^102–105^. To confirm the accuracy of our cellular measurements, we calculated the dimensions of microspheres (LIVE/DEAD BacLight Bacterial Viability and Counting Kit, ThermoFisher Scientific) with a reported diameter of 6 µm and obtained an average diameter of 5.94 ± 0.07 µm (N=20).

### Grams per dry cell weight to OD_600_ measurements

To perform gDCW measurements, three biological replicates of each microbe were grown as previously described, 500 mL (*Z. mobilis* and *E. coli*) or 100 mL (*C. thermocellum*) of bacterial culture was collected during late-phase growth (OD_600_ > 0.75), and the culture was centrifuged at 4,255 × g for 20 mins at 4 °C. Cell pellets were washed with ddH_2_O to remove salts, and this cell suspension was vacuum filtrated through a pre-weighed 0.45-µm-pore-size hydrophilic nylon filter (Millipore catalog no. HNWP04700) applied to a sintered glass funnel. The nylon filter containing cells was placed in a glass petri dish and oven dried at 80 °C until the mass of the filter was stable (24-48 hours)^30,78^. The mass of the cells on the filter was then normalized to the OD_600_ at the time of collection and the culture volume that was filtered to obtain gDCW OD_600_^−1^ L^−1^ (Table S2).

### Sugar uptake and growth rate calculations

Growth rates (hr^−1^) and sugar (glucose: *Z. mobilis* and *E. coli*; cellobiose: *C. thermocellum*) consumption rates (mmol_sugar_ gDCW^−1^ hr^−1^) were obtained by growing three biological replicates of each microbe as previously described. OD_600_ measurements and 1 mL culture samples were collected every hour until stationary phase was achieved. Culture samples were centrifuged at 21,000 × g for 5 min at 4 °C, and the supernatant was stored at −80 °C until analysis by LC-MS. Samples were diluted 1:100 (*Z. mobilis*) or 1:20 (*E. coli* and *C. thermocellum*) with HPLC-grade H_2_O, mixed 50:50 with 1 mM [U-^13^C] glucose (*Z. mobilis* and *E. coli*) or 1 mM [U-^13^C] cellobiose (*C. thermocellum*), and analyzed via LC-MS. LC-MS analysis was performed on a Vanquish ultra-high-performance liquid chromatography (UHPLC) system (Thermo Scientific) coupled to a hybrid quadrupole-Orbitrap mass spectrometer (Q Exactive; Thermo Scientific) equipped with electrospray ionization operating in negative-ion mode. The chromatography was performed at 25°C using a 2.1-by 100-mm reverse-phase C_18_ column with a 1.7-μm particle size (Water; Acquity UHPLC ethylene-bridged hybrid). The chromatography gradient used Solvent A (97:3 H_2_O:methanol with 10 mM tributylamine adjusted to pH 8.2 using 10 mM acetic acid) and Solvent B (100% methanol) and was as follows: 0–2.5 min, 5% B; 2.5–8 mins, linear gradient from 5% B to 95% B; 8–10.5 min, 95% B; 10.5–11 min, linear gradient from 95% B to 5% B; 11– 15 min, 5% B. The flow rate was held constant at 0.2 mL min^−1^. The MS parameters used were as follows: full MS-single ion monitoring (SIM) scanning between 70 and 1,000 *m/z*; automatic gain control (AGC) target, 1e6; maximum injection time (IT), 40 ms; resolution of 70,000 full width at half maximum (FWHM). Data analysis was performed using the MAVEN software^106^. Glucose and cellobiose were identified based on retention times matched to pure standards. The ratio of ^12^C-to-^13^C peak intensities was used to calculate glucose or cellobiose concentrations and sugar consumption rates were normalized to gDCW and growth rates (Table S2).

### Cell enumeration via flow cytometry

Cell densities (cell mL^−1^) were quantified using flow cytometry. Three biological replicates of each microbe were grown as previously described. When cells reached an OD_600_ of 0.45, 5 mL of bacterial culture was collected and centrifuged at 4,255 × g for 5 mins at 4 °C. Cell pellets were washed twice with NaCl solutions to remove media components. NaCl solutions were prepared at 0.85, 0.55, and 0.27% to match the osmolarity of M9, ZMM, or MTC media, respectively, to prevent cell lysis/ plasmolysis. Cells were then diluted 1:100 in NaCl solution, equimolar amounts of SYTO 9 and propidium iodide, and 10^6^ counting beads (LIVE/DEAD BacLight Bacterial Viability and Counting Kit, ThermoFisher Scientific). Samples were then immediately analyzed via flow cytometry.

Prior to acquisition, sample tubes were briefly vortexed. Samples were analyzed using an Attune NxT Acoustic Focusing Cytometer (ThermoFisher Scientific) using the following settings: flow rate, 12.5 μL min^−1^; FSC-A, 300; SSC-A, 325; BL1-A, 350; YL2-A, 500; RL3-A, 400; VL1-A, 400. For each replicate, 50 μL equating to approximately 100,000 single cells were analyzed/ counted. Data analysis was performed using the FlowJo software (BD Biosciences, version 10.9). Manual gating using an FSC-A vs SSC-A dotplot was performed to distinguish cells and beads from debris and aggregates, and an SSC-A vs SSC-H dotplot was used to account for smaller aggregates and multiplets. Cell numbers were then calculated using the following formula: # *of bacterial events* × *dilution factor*/# *of bead events* × 10^6^ (Table S2).

### Protein cost calculations and in vivo flux and thermodynamic data

*In vivo* free energies and glycolytic fluxes were obtained from previous studies that quantified these values via MFA models (Table S1 and S9). These flux and thermodynamic data were calculated under similar growth conditions used in this study^30–32^. Intracellular fluxes and free energies under N_2_ fixation conditions in *Z. mobilis* were also obtained from previous MFA data^91,92^. To quantify protein costs, we normalized the sum of all participating enzymes and isoenzymes to the intracellular flux of the metabolic reaction. For example, the protein cost for the PFK reaction in *E. coli* equates to the total concentration of the isoenzymes PfkA and PfkB (260.9 μg gDCW^−1^) normalized to the *in vivo* flux (8.23 mmol/ (gDCW hr^−1^)). To quantify the total protein cost of fermentation in *Z. mobilis*, we took the ratio of the sum of Pdc, Adh (AdhA and AdhB), AldB, and Ldh (Ldh1 and Ldh2) enzyme concentrations (μg gDCW^−1^) to the combined flux of Adh and AldB. Lactate flux data was unavailable but is largely considered to be negligible in *Z. mobilis*. The protein cost of just ethanol fermentation in *Z. mobilis* was determined first by normalizing the Pdc enzyme concentration to the ratio of acetate to Pdc flux, which provided the proportion of Pdc enzyme strictly dedicated to ethanol production. The ethanol fermentation protein cost was then quantified by taking the ratio of the sum of the adjusted Pdc enzyme concentration and Adh enzyme concentration to the Adh flux.

To quantify the total protein cost of fermentation in *C. thermocellum*, we first quantified the proportion of Pfor and Pfl (i.e., the sum of all Pfor subunits, Pfl, and Pfl-activating enzyme, see Table S11) protein dedicated towards fermentation metabolites (i.e., acetate and ethanol). This was done by calculating the ratio of acetate and ethanol (i.e., Adh) flux to the total acetyl-CoA flux (i.e., the sum of Pfor and Pfl flux)^107^. This ratio represented the proportion of Pfor and Pfl enzyme used for fermentation. These normalized Pfor and Pfl protein concentrations were combined with the protein concentrations for Pta, Ack, Ldh, and Adh (Aldh/ Adh, Adh1-5), and this total protein sum was normalized to the sum of lactate, acetate, and ethanol flux. The protein cost of just ethanol fermentation in *C. thermocellum* was determined first by normalizing the Pfor and Pfl enzyme concentrations to the ratio of ethanol flux to the total acetyl-CoA flux, which provided the proportion of Pfor and Pfl enzyme strictly dedicated to ethanol production. The sum of these adjusted Pfor and Pfl protein levels and total Adh enzyme concentration was then normalized to the ethanol flux. All fluxes obtained from the literature were normalized to our glucose uptake rates (Tables S2 and S9).

To calculate free energies for the *E. coli* PTS and the *C. thermocellum* reactions cellobiose phosphorylase, phosphoglucomutase, glucokinase, and pyruvate phosphate dikinase/ malate shunt that lack intracellular data, we combined *in vivo* metabolite concentration data^30,31,108–110^ with standard Gibbs free energy estimates^81^ and obtained theoretically optimized free energies for these reactions using the Max-Min driving force (MDF) computational tool (Table S1). The MDF method identifies the most thermodynamically restrictive reactions in a pathway and maximizes their thermodynamic driving force by optimizing metabolite concentrations^16^. MDF analysis was performed using the Python package equilibrator-pathway (version 0.5.0)^111^. Intracellular pH, pMg, and ionic strength were set to 7, 3, and 250 mM, respectively. Temperature was set to 310.15 K and 328.15 K for *E. coli* and *C. thermocellum*, respectively. Maximum and minimum metabolite concentration bounds were based on a 50% range of absolute intracellular data^30,31^. Importantly, these *in vivo* metabolite concentrations were quantified in *E. coli* RL3000 and *C. thermocellum* DSM1313 cells grown under equivalent conditions used in this study. Cellobiose and glucose concentration bounds were informed by cellobiose and glucose concentrations in the media at the time of protein quantification (i.e., OD_600_ 0.45). Diphosphate concentration bounds for *C. thermocellum* were based on intracellular concentration data quantified in related *Clostridia* species that encode for PPi-PFKs^109,110^. Orthophosphate concentration bounds for *E. coli* were based on measurements performed in *E. coli* K-12^108^. For both microbes, the minimum bound of pyruvate was increased from the default 1 μM to 1 mM based on the intracellular concentrations of other glycolytic intermediates. SBtab files used to perform the MDF analyses for *E. coli* and *C. thermocellum* can be found in Tables S17 and S18, respectively.

### COG classification of proteins

Proteins were assigned to COG-defined cellular functions using the National Center for Biotechnology Information (NCBI) Batch CD-Search tool^82,83,112,113^. Searches were performed against the COG database. Unassigned proteins were manually classified with the ‘Function unknown’ COG category. The percentage of the proteome mass dedicated to each COG-defined cellular function was quantified on a mass basis (fg cell^−1^). For proteins with multiple COG classifications, the protein concentration was evenly divided amongst each category.

### Isotopically labeled ethanol and acetate experiments

To assess the reversibility of the fermentation pathways in *Z. mobilis* and *C. thermocellum*, we performed growth experiments with isotopically labeled ethanol or acetate and tracked the propagation of isotope labeling to upstream intermediates. *Z. mobilis* was grown as previously described. When cells reached an OD_600_ of 0.5, 7.5 mL of bacterial culture was collected for four biological replicates. Cells were centrifuged, the supernatant was discarded, and two replicates of cell pellets were resuspended in either 7.5 mL of fresh ZMM spiked with 2.5 g L^−1^ of 1-^13^C-ethanol or 1-^13^C-acetate. Cells were grown for an additional 45 minutes before metabolites were extracted. *C. thermocellum* was also grown as previously described. Two biological replicates were grown in either MTC media prepared with 2 g L^−1^ of 2-^13^C-ethanol or 1,2-^13^C-acetate. Metabolite extractions were performed when cells reached an OD_600_ of 0.45.

At the time of metabolite extraction, 5 mL of liquid culture was collected in the anaerobic chamber using a serological pipette. Cells were separated from the media by vacuum filtering the culture through a 0.45-μm-pore-size hydrophilic nylon filter (Millipore; catalog no. HNWP04700) applied to a sintered glass funnel. The nylon filter containing cells was immediately immersed cell-side down into a plastic petri dish (5.5-cm diameter) containing 1.5 mL cold (–20°C) extraction solvent (40:40:20 by % volume methanol-acetonitrile-water; all high-performance liquid chromatography [HPLC] grade) and kept on a chilled aluminum block. This process simultaneously lysed the cells, quenched metabolism, and dissolved intracellular metabolites. The petri dish was lightly swirled to ensure complete contact of solvent with the filter. Filters remained in the cold solvent for ∼15 min before being repeatedly rinsed in the extraction solvent to collect any remaining cell debris and metabolites. The cell-solvent mixture was then transferred to a 1.5-mL microcentrifuge tube, removed from the anaerobic chamber, and centrifuged at 16,000 × *g* for 10 min at 4°C, and the supernatant was collected for LC-MS analysis.

LC-MS analysis was performed as previously described but with altered chromatography. The chromatography gradient used Solvent A and Solvent B and was as follows: 0 to 2.5 min, 5% B; 2.5 to 17 min, linear gradient from 5% B to 95% B; 17 to 19.5 min, 95% B; 19.5 to 20 min, linear gradient from 95% B to 5% B; and 20 to 25 min, 5% B. Data analysis was performed using the MAVEN software^106^. Pyruvate and acetyl-CoA were identified based on retention times matched to pure standards. Metabolite mass isotopomer distributions from ^13^C labeling samples were corrected for ^13^C natural abundance using ElemCor^114^. Pyruvate labeling patterns were calculated from valine to exclude unlabeled (M+0) pyruvate in the media, and the acetyl group labeling in acetyl-CoA was calculated from aspartate and glutamate.

## Data availability

All raw proteomics data will be made available on request. Bacterial strains will be made available upon request.

## Author contributions

D.A. and D.B.K wrote the main manuscript text. D.A. and D.B.K prepared figures and tables. D.B.K., A.J., E.S., E.T., J.W., A.H., D.M.S. performed experiments and analyzed data. A.J., E.S., and J.C. designed and performed the proteomics mass spectrometry experiments. All authors reviewed the manuscript.

## Disclosure and competing interests statement

J.J.C. is on the Scientific Advisory Board of Seer. J.J.C. is a consultant for Thermo Fisher Scientific and a founder of CeleramAb Inc.

## Acknowledgements

Funding provided by The Center for Bioenergy Innovation (under Award Number ERKP886) and the Great Lakes Bioenergy Research Center (under award no. DE-SC0018409), both U.S. Department of Energy Research Centers supported by the Office of Biological and Environmental Research in the DOE Office of Science. This work was also funded by the DOE Early Career Research Program under award no. DE-SC0018998 (to D.A.). Additional support was provided by the National Institutes of Health grant R35 GM118110 (to J.J.C.).

---

**Figure S1.**
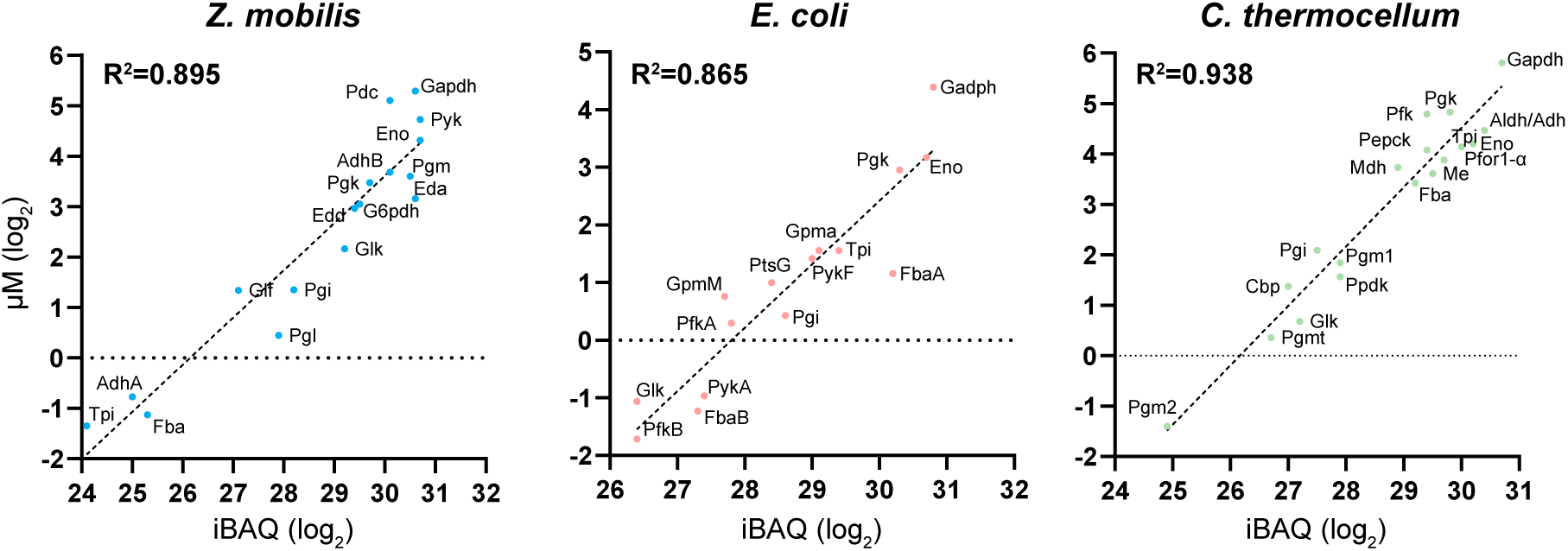
Correlation between AQUA quantitation and iBAQ values. Linear regression analyses between log_2_ transformed absolute quantification (AQUA) values (y-axis) versus intensity-based absolute quantification (iBAQ) values (x-axis) for each bacterium. Pearson’s square correlation coefficients (R^2^) are displayed on all plots. Abbreviations: glucose facilitated diffusion protein (Glf), glucokinase (Glk), glucose 6-phosphate dehydrogenase (G6pdh), phosphogluconolactonase (Pgl), 6-phosphogluconate dehydratase (Edd), phosphoglucose isomerase (Pgi), 2-keto-3-deoxy-6-phosphogluconate aldolase (Eda), fructose 1,6-bisphosphate aldolase (Fba, FbaA, FbaB), triose phosphate isomerase (Tpi), glyceraldehyde 3-phosphate dehydrogenase (Gapdh), phosphoglycerate kinase (Pgk), phosphoglycerate mutase (Pgm, GpmM, GpmA, PGM1, PGM2), enolase (Eno), pyruvate kinase (Pyk, PykA, PykF), pyruvate decarboxylase (Pdc), alcohol dehydrogenase (AdhA, AdhB), PTS system glucose-specific EIICB component (PtsG), phosphofructokinase (PfkA, PfkB, Pfk), cellobiose phosphorylase (Cbp), phosphoglucomutase (Pgmt), pyruvate phosphate dikinase (Ppdk), phosphoenolpyruvate carboxykinase (Pepck), malate dehydrogenase (Mdh), malic enzyme (Me), pyruvate ferredoxin oxidoreductase I alpha domain (PFOR1-α), bifunctional acetaldehyde/alcohol dehydrogenase (Aldh/ Adh).

**Figure S2.**
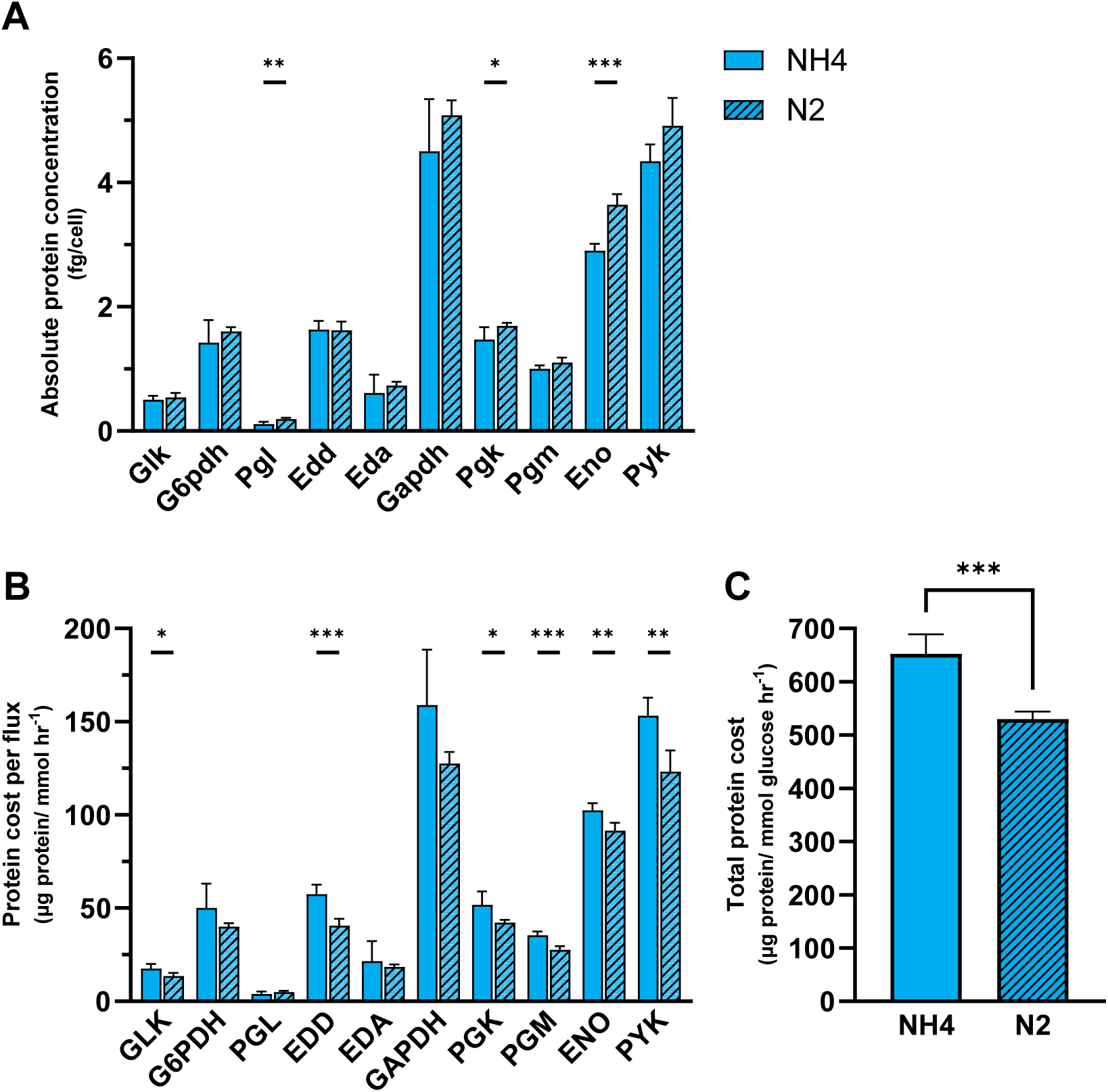
Protein cost comparisons for the ED pathway in *Z. mobilis* grown under N_2_-fixing vs. NH_4_^+^-replete conditions. **A.** Absolute concentrations of glycolytic enzymes (fg/cell^−1^) in *Z. mobilis* grown with NH_4_^+^ (solid bars) or N_2_ (striped bars) as the sole nitrogen source. Enzyme concentrations for *Z. mobilis* cells grown on N_2_ were quantified by normalizing previous shotgun proteomics data to absolute values (Materials and Methods)^92^. **B.** Comparisons of protein costs for ED pathway reactions (µg protein/ (mmol hr^−1^)) between NH_4_^+^-replete conditions and N_2_-fixing conditions. Protein costs for each glycolytic reaction were calculated by normalizing the sum of all participating enzymes to the intracellular flux of the reaction. **C.** Total protein cost of the ED pathway in *Z. mobilis* grown on NH_4_^+^ vs. N_2_. The total protein cost for the ED pathway in each condition represents the sum of all glycolytic enzymes normalized to the glucose uptake rate of *Z. mobilis* grown under that condition (Materials and Methods). Data for each nitrogen condition represent the averages of five biological replicates. Error bars show ± standard deviation. Some error bars are too small to be visible in this representation. Asterisks indicate statistical significance: *, *P* < 0.05; **, *P* < 0.01; ***, *P* < 0.001. See Table S13 for absolute enzyme concentrations under N_2_ conditions. Abbreviations: glucokinase (GLK), glucose 6-phosphate dehydrogenase (G6PDH), phosphogluconolactonase (PGL), 6-phosphogluconate dehydratase (EDD), 2-keto-3-deoxy-6-phosphogluconate aldolase (EDA), glyceraldehyde 3-phosphate dehydrogenase (GAPDH), phosphoglycerate kinase (PGK), phosphoglycerate mutase (PGM), enolase (ENO), pyruvate kinase (PYK)

## Notes

### Competing Interest Statement

The authors have declared no competing interest.

